# Extracellular vesicles from antigen-activated B cells stimulate oncogenic herpesvirus lytic gene expression

**DOI:** 10.64898/2026.02.09.704805

**Authors:** Amna Music, Hedvig Djupenström, Diogo M Cunha, Saara Hämälistö, VTR Sikha, Milla Majaniemi, Knut KB Clemmensen, Marja Jäättelä, Kenji Maeda, Silvia Gramolelli, Pieta K. Mattila

## Abstract

B cell-derived extracellular vesicles (B-EVs) have been associated with immunomodulatory functions, including antigen presentation, cytokine signaling, and T cell activation. However, little is known about the general composition of B-EVs or how the state of the donor cells affects their composition or functions. We performed integrated proteomic, lipidomic, and transcriptomic analyses of EVs secreted by B cells under resting conditions or after 30 min of B cell receptor (BCR) activation. Antigen-driven BCR stimulation led to acute remodeling of the protein and lipid composition, and the RNA cargo of the B-EVs, which also translated into differential responses in recipient B cells. Remarkably, EVs from activated B cells stimulated lytic gene expression of latent oncogenic γ-herpesviruses Epstein–Barr virus (EBV) and Kaposi sarcoma-associated herpesvirus (KSHV). These findings suggest a novel mechanism for the potentially oncogenic reactivation of these lifelong-persisting latent herpesviruses, residing in B cells. Our results support a mechanism in which, upon immune activation, B-EVs from antigen-specific B cells induce viral reactivation in bystander B cells in a paracrine manner, providing molecular insights that bridge immune activation and the pathogenesis of γ−herpesvirus infections.

## Introduction

Extracellular vesicles (EVs) have emerged as a powerful means of intercellular communication. Secreted by all cell types via endosomal pathways (exosomes) or plasma membrane budding, EVs carry diverse bioactive molecules, including enzymes, regulatory RNAs, and lipids, derived from the host cells. EVs can modulate recipient cells and the tissue microenvironment both locally and systemically^1–3^. Their heterogeneity and biocompatibility endow EVs with multifaceted functions and applications, including regulation of homeostasis, immune responses, and cancer progression, as well as use as biomarkers and drug delivery carriers^4–7^.

Within the immune system, EVs serve as important signaling messengers carrying antigens, cytokines, and regulatory molecules. They have been reported to influence responses ranging from antimicrobial and antitumor defenses to immunological tolerance and inflammation^8,9^. For example, activation of T cells remodels their EV composition and release, with notable effects on antigen-presenting cells (APCs) and their effector responses^10–13^. These findings highlight EVs as integral players in the immune response and emphasize their functional heterogeneity depending on cellular activation state.

B cells, beyond their well-established role in antibody production also function as APCs, secrete cytokines and release EVs (B cell-derived EVs; B-EVs). B-EVs carry major histocompatibility complexes I and II (MHC-I, MHC-II), tetraspanins, and the B cell antigen receptor (BCR)^14–17^. Early work in B cell lines and stimulated primary cells has shown that B-EVs can present peptide-antigens on the MHC-II and shape T cell responses similarly to antigen-presenting cells (APCs)^14,18,19^. Various stimulations on B cells have also been shown to translate into differences in B-EV quantity and quality. For example, stimulation via CD24 and BCR increases EV production in murine B cells and promotes transfer of functional BCR to recipient cells^20^. Exosome release can also be triggered via innate immune stimulation by CD40 and IL-4 in primary B cells^21^. However, it is unknown whether this release is due to immediate receptor-mediated cellular signaling or to long-term changes in cell state and differentiation. BCR engagement rapidly triggers extensive cell signaling leading to acute cellular changes like plasma membrane remodeling as well as transcriptional reprogramming^22–24^, suggesting that also B-EVs could undergo dynamic changes at several stages after BCR activation. A comprehensive comparison of EVs from resting and antigen-activated B cells is currently lacking, and it remains unclear how acute antigen engagement reshapes the composition and immunological properties of B-EVs.

B cells are natural hosts of oncogenic γ-herpesviruses: Epstein–Barr virus (EBV) and Kaposi sarcoma-associated herpesvirus (KSHV). Over 95% of the world population carries latent EBV in their B cells, whereas KSHV is prevalent in sub-Saharan Africa and the Mediterranean basin^25,26^. These viruses have been identified as the causal agents of a broad range of disorders, yet only a fraction of infected subjects experience pathogenic symptoms. Herpesviruses establish lifelong infections in the human host, and to date, neither specific therapy for eradication nor vaccines to prevent infections exist for EBV and KSHV^25,26^. While these infections are predominantly latent, characterized by limited gene expression and persistence of the viral genome as a non-integrated episome, sporadic lytic reactivation drives viral propagation and has been linked to disease pathogenesis^27–29^. The lytic cascade is initiated by key viral transcription factors encoded by KSHV ORF50 and EBV BZLF1, which drive the temporally orchestrated expression of downstream lytic genes, culminating in the assembly and release of progeny viral particles^30^. Although the physiological stimuli driving herpesvirus lytic expression are still poorly understood, several studies have indicated that autocrine signaling from activated B cells through the BCR pathway can induce lytic reactivation^31–34^. Whether secreted B-EVs could mediate a paracrine effect and induce viral reactivation in neighboring cells has, however, not yet been investigated.

Here, we provide an integrated characterization of the proteomic, lipidomic, and transcriptomic profiles of EVs released by resting and acutely activated B cells, combined with analysis of recipient cells carrying latent KSHV and EBV-infections. We demonstrate that antigenic activation of B cells leads to a rapid modulation of the secreted EVs, equipping them with the capacity to increase lytic gene expression in recipient cells. Our findings uncover a previously unrecognized route of B cell-mediated communication and provide novel insight into the regulation of host-virus dynamics.

## Results

### Characterization of B-EVs secreted from resting and activated B cells

To investigate the possible antigen-induced changes in EVs secreted by B cells, we first set up the purification protocol for B-EVs from two cultured human B cell lines, Raji D1.3 and BJAB (Fig. 1A). Antigen engagement in B cells triggers an immediate and strong, multibranched intracellular response. To investigate the possible acute effects of the BCR stimulation on the secreted B-EVs, they were collected from the supernatant of either non-activated, resting state cells or after 30-minute activation with surrogate antigen engaging the BCR. F(ab’)_2_ fragments of anti-human IgM antibodies (αIgM) were used as surrogate antigen for BJAB cells, matching their endogenous BCR, while F(ab’)_2_ fragments of anti-mouse IgM antibodies were used for Raji D1.3 cells, which carry a transgenic IgM BCR of mouse origin. EV-enriched samples were obtained by size exclusion chromatography (SEC) from B cell culture medium, concentrated and subjected for all downstream analyses following the MISEV guidelines^35^. These samples are referred to as B-EVs throughout this manuscript. The presence of EVs in our preparations was verified by transmission electron microscopy (TEM), where vesicular structures consistent with the morphology of EVs were observed (Fig. 1B, C). We also observed protein aggregates, often adhered to the vesicles, in the TEM images, indicating that our EV preparations also contain some secreted non-vesicular proteinaceous material or other extracellular particles (Fig. S1A, B), which are often found in EV isolations^35,36^. Nanoparticle Tracking Analysis (NTA) indicated a shift in EV size profile upon cell activation, particularly in Raji D1.3 cells (Fig. 1D, S1C); however, NTA showed notable fluctuations between the replicates (Fig. S1C). The protein quantification measurements showed significantly higher protein concentration in EV samples from antigen-activated Raji D1.3 cells (Fig. 1E). A similar trend appeared also appeared in BJAB cells, however, with no statistical significance. Variation in the EV yield between different preparations was also clear on the protein quantification. We confirmed the presence of established EV protein markers Ezrin, Alix and CD81, in our preparations by Western blot analysis (Fig. 1F, S1D). Lack of Calnexin indicated the absence of cellular contamination in our isolated EV samples. Together, these analyses provided us with confidence in our EV preparation protocol and enabled us to proceed with downstream analyses.

**Figure 1.**
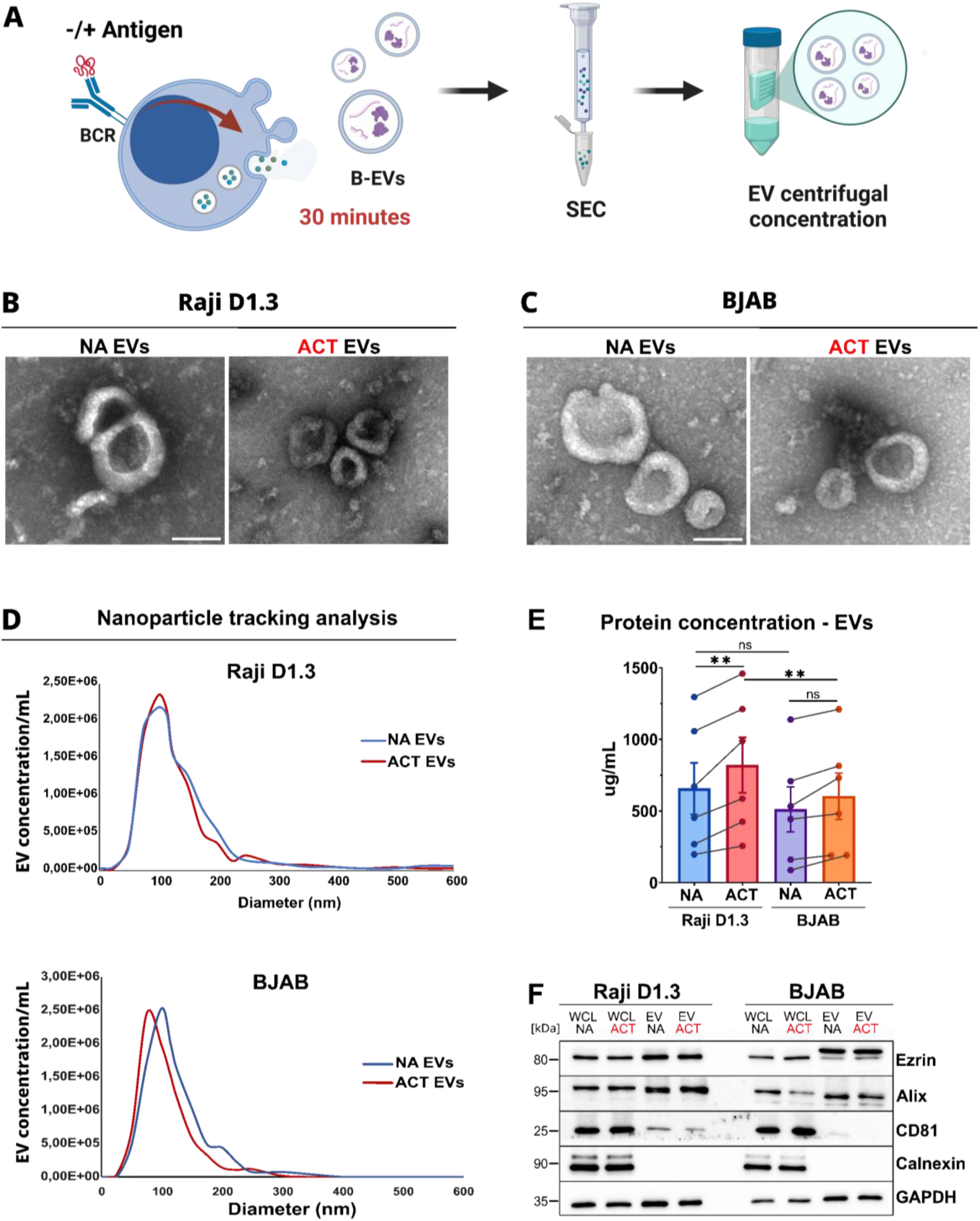
Isolation and characterization of B-EVs from resting and antigen-activated B cell lines. **(A)** Schematic illustration of B-EV isolation workflow. **(B-C)** SEC-isolated and pooled B-EV samples from Raji D1.3 (B) and BJAB (C) cells, either left non-activated (NA) or activated with surrogate antigen (αIgM F(ab’)_2_, 10 µg/mL) for 30 minutes (ACT), and imaged by TEM; scale bar: 100 nm. **(D)** NTA analysis of the EV size distribution between NA and ACT samples prepared as above. Representative graphs shown, n = 4 (see also the supplementary Figure S1C). **(E)** BCA protein assay to measure the protein concentrations of NA and ACT samples prepared as above. Data is presented as mean ± SEM, n=6. Statistical significance was determined using two-tailed paired Student’s T test *p < 0.05, **p < 0.01, ***p < 0.001. **(F)** Western blot analysis of Raji D1.3 and BJAB cell lysates and their corresponding B-EV samples. Equal amounts of protein were loaded to evaluate EV enrichment relative to whole-cell lysates (WC). EV markers CD81, Alix, and Ezrin were used to confirm EV identity, GAPDH served as a loading control, and Calnexin was included as a negative control for cellular contamination.

### Lipidomic analysis suggests the plasma membrane as the major source of B-EVs

To gain insight into the mechanisms by which antigenic exposure modulates B-EV secretion, we next investigated the lipid composition of B-EVs secreted by resting or activated B cells using shotgun lipidomics^37^. When compared to corresponding whole cell (WCs) lysates, this method can suggest the organelle of origin of the EVs.

In total, we identified 489 lipid species spanning glycerolipids (GLs), glycerophospholipids (GPLs), sphingolipids (SLs), and sterol lipids (STs), across 29 lipid classes (Supplementary Table SI). Absolute quantification was achieved using internal standards, and values were normalized to mol% of total lipids. Cholesterol was excluded from the presented dataset due to variability across independently performed experiments, likely caused by its poor ionization in the instrument.

Heatmap visualization of mol% values revealed distinct lipid profiles between B-EVs and WCs, and between Raji D1.3 and BJAB cells, with high consistency across the three biological replicates (Fig. 2A). To explore enrichment or depletion of given lipids in B-EVs, we calculated log₂-transformed fold changes (B-EV/WC) in mol% for each species (Fig. 2B). In parallel, we assessed the impact of cellular activation by comparing lipid profiles of B-EVs from activated and resting cells (Fig. 2C). Although the precise enrichment patterns varied between the two cell lines, comparison of log₂ fold-change profiles revealed positive Spearman correlation coefficients between Raji D1.3 and BJAB cells (Fig. 2D), indicating that the overall patterns of lipid enrichment into B-EVs were conserved across cell types.

**Figure 2.**
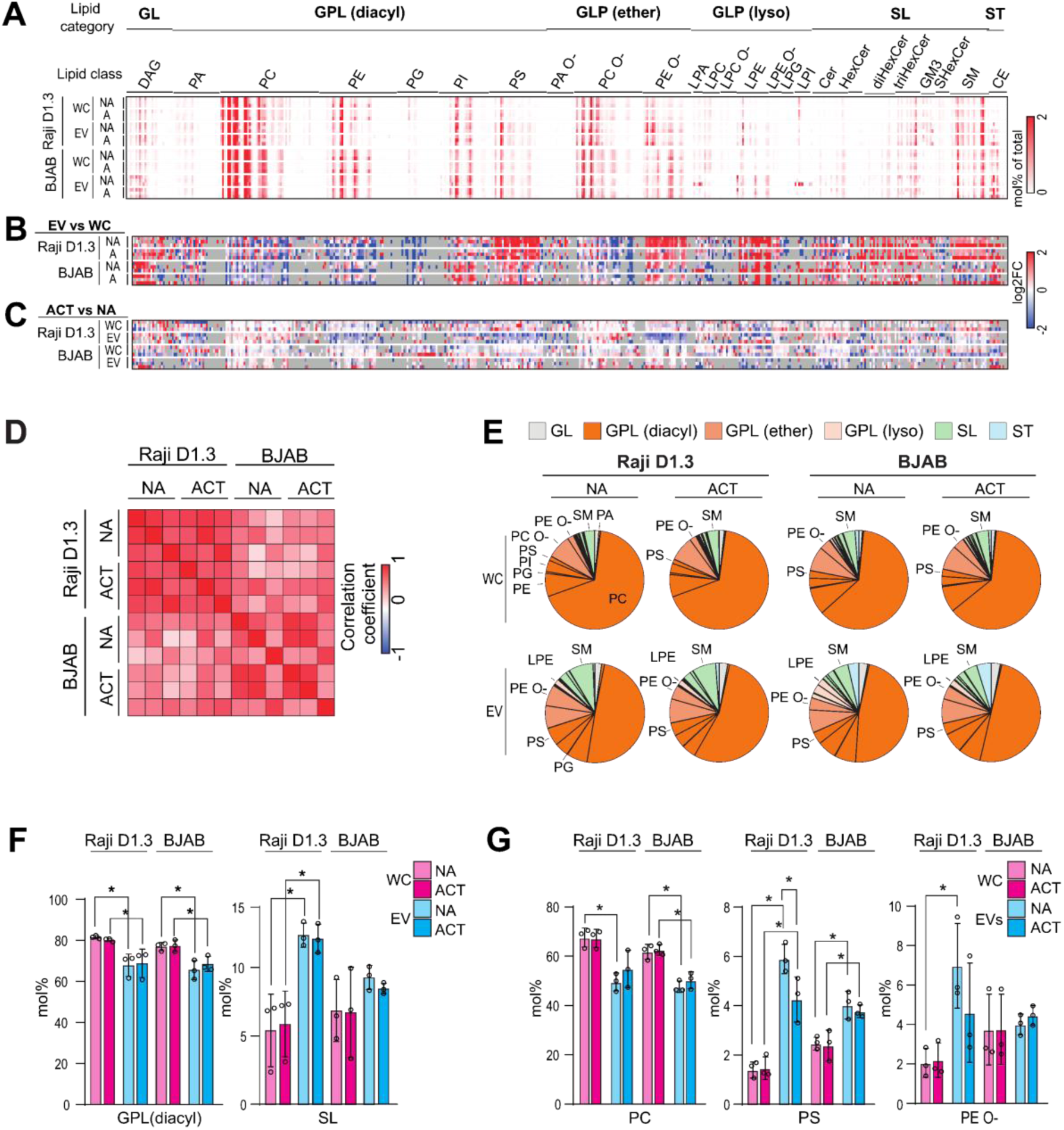
Shotgun lipidomics analysis exposes plasma membrane-like lipid composition in B-EVs. **(A)** Heatmap showing mol% values of lipid species identified in EVs and whole cell (WC) lysates and used in comparative analyses. Three independently performed experiments were performed for 30 minutes activated (ACT) and non-activated (NA) cells, as previously described. **(B-C)** Heatmaps showing log2-transformed fold changes of lipid species between the EVs and WCs (EV/WC) (B) or between the activated and non-activated (ACT/NA) B-EVs and WC lysates (C). **(D)** Correlation matrix displaying Spearman correlation coefficients between cell lines and conditions (activated or non-activated) calculated from the log2-transformed fold changes (EV/WC). **(E)** Pie charts displaying the mol% of lipid classes and categories. **(F)** Bar graphs displaying mol% of total SL and total diacyl GPLs. **(G)** Bar graphs showing mol% of the O-classes of PC, PS, and PE. * indicates P < 0.05 in t-test. N=3

Our analysis comparing B-EV and whole cell lipidomes revealed that B-EVs—especially those from Raji D1.3 cells—showed consistent enrichment of SLs, with broad upregulation across many SL species, in contrast to glycerophospholipids (GPLs), where species in some classes were enriched, and others depleted. Consistent with the widespread enrichment of SL species (Fig. 2B), B-EVs displayed a higher total SL content and a corresponding reduction in diacyl GPLs compared to their respective WC lipidomes (Fig. 2E, F). Among the GPLs enriched in B-EVs were phosphatidylserine (PS), ether-linked phosphatidylethanolamine (PE O-), and lysophosphatidylethanolamine (LPE). Notably, SLs and PS are typically enriched at the plasma membrane^38^ and we have previously found PE O- and LPE to be enriched in plasma membrane fractions from HeLa cells using similar shotgun lipidomics^39^.These findings suggest that a significant fraction of B-EVs would be derived from the plasma membrane. Notably, we did not observe enrichment of bis(monoacylglycero)phosphate (BMP) in B-EVs. BMP species are structural isomers of phosphatidylglycerol (PG) and are therefore reported as PG in our shotgun lipidomics dataset. As BMP is characteristically localized to late endosomes and lysosomes, its lack of enrichment in B-EVs is consistent with a limited contribution from multivesicular body–derived membranes to the B-EV lipidome.

Upon cell activation, the lipidomic changes were largely confined to B-EVs from Raji D1.3 cells. Activation consistently led to reduced levels or showed trends toward depletion of PS and PE O– species (Fig. 2C), as well as their corresponding classes (Fig. 2G). In contrast, other major lipid classes—including phosphatidylcholine (PC) and SLs—remained largely unchanged (Fig. 2C, F, G). No marked changes were observed in B-EVs from BJAB cells or in the whole-cell lipidomes from either cell line, suggesting that the observed changes reflect selective lipid remodeling in Raji D1.3-derived B-EVs rather than global shifts in cellular lipid metabolism. The shift detected in Raji D1.3-derived B-EVs suggests some alterations in the plasma membrane lipid profile, which could be linked to the strong signaling activity ongoing at the plasma membrane upon BCR activation, that also includes the stabilization of lipid raft-like domains and formation of lipid-linked second messengers, such as diacylglycerol and inositol triphosphate^40,41^.

Taken together, the lipidomic analyses suggested that the B-EVs were largely originated by plasma membrane budding. Although antigenic activation of donor cells led to some changes in the lipid profile of B-EVs, the plasma membrane character remained, indicating that the activation-induced lipidomic changes originate rather from modulation of the plasma membrane than from changes in the cellular source per se.

### Proteomic and transcriptomic analyses of B-EVs show distinct changes upon acute antigen-mediated cell activation

To further characterize the molecular landscape of B-EVs and the changes upon antigen activation, we performed label-free proteomics on SEC-isolated EVs derived from Raji D1.3 and BJAB cells activated or not for 30 min with surrogate antigen, and matching WC samples. Overall, parental cells and B-EVs showed markedly different global proteomic profiles. B-EV samples exhibited greater technical variance between replicates than the variance induced by antigen engagement, denoting the heterogeneity between the B-EV preparations from different batches of cells and the relative subtlety of the changes in protein expression induced by cell activation (Fig. 3A, S2A).

**Figure 3.**
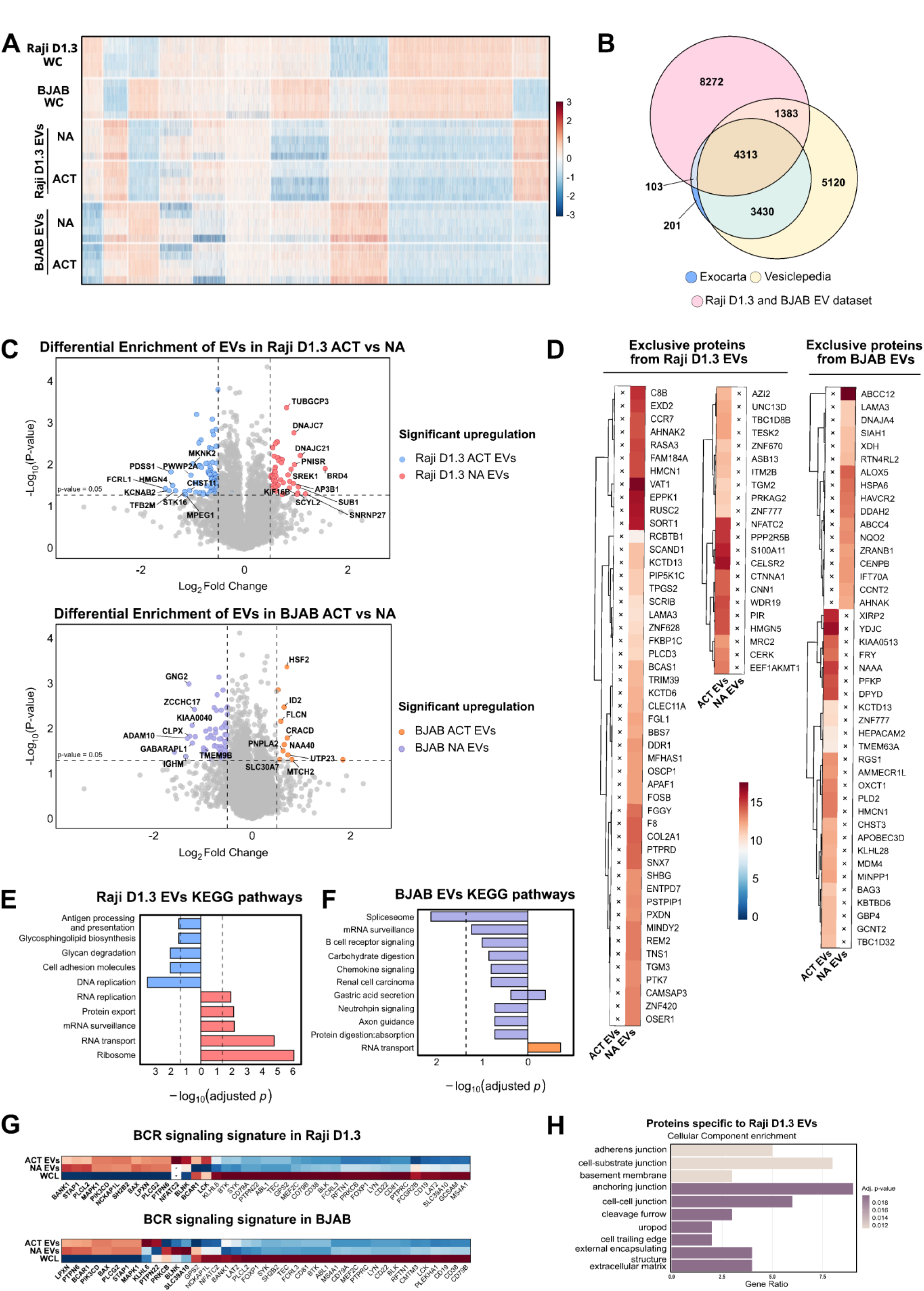
Proteomic profile of the B-EVs is altered by acute antigen activation. **(A)** Heatmap of protein expression profiles across cell lines. Protein intensities were normalized with z-score transformation and B-EVs isolated from 30 minutes (αIgM F(ab’)_2_) activated (ACT) or non-activated (NA) conditions were compared to whole cell lysates (WC) of both cell lines. Proteins were clustered based on hierarchical clustering based on Ward method. Color scale: Red indicates higher expression relative to the mean, blue indicates lower expression (z-score scale from -3 to +3). **(B)** Venn diagram of proteins identified in the combined B-EVs from both cell lines with proteins annotated in the Exocarta and Vesiclepedia databases. **(C)** Volcano plots of differentially expressed (DE) proteins in BJAB and Raji D1.3 B-EVs. The plot displays log2 fold change (FC) (x-axis) versus -log10 p-value (y-axis) for activated (ACT) conditions compared to resting (NA). Colored points (right; red and orange) represent significantly upregulated proteins in ACT B-EVs (FC > 0.5, p < 0.05), colored points (left; blue and purple) show significantly upregulated proteins in NA B-EVs (FC < -0.5, p < 0.05), and grey points are non-significant. Dashed lines indicate significance thresholds. Top 10 most significantly up/down-regulated proteins are labelled. **(D)** Heatmap of proteins exclusively identified in the B-EVs and not in WCL, in Raji D1.3 and BJAB. Protein abundance (log2 intensity) is displayed. Color scale reflects relative protein abundance. Conditions highlighted by a cross (X) represent the absence of the protein. **(E, F)** KEGG pathway enrichment analysis of BJAB and Raji D1.3 B-EVs. Mirrored bar plot shows enriched pathways (BH-corrected) in B-EVs from activated (red for Raji D1.3 and orange for BJAB) and resting cells (blue for Raji D1.3 and purple for BJAB). X-axis displays -log10 (adjusted p-values), with pathways ordered by significance. The dashed line indicates the significance threshold (adjusted p-value < 0.05). **(G)** Heatmap with BCR signaling-associated proteins in BJAB and Raji D1.3. Protein abundance was z-score normalized across ACT B-EVs, NA B-EVs and WCL conditions. Red indicates higher expression relative to the mean, blue indicates lower expression (z-score scale from -1 to +1). Proteins were matched against BCR signaling pathway genes (GO:0050853) for BCR associated genes. Conditions highlighted by a cross (X) represent the absence of the protein. **(H)** GO:CC enrichment analysis of proteins exclusively identified in Raji D1.3-derived B-EVs. Enrichment was performed using clusterProfiler with all detected proteins as the background universe and Benjamini–Hochberg correction. The bar plot shows the top enriched cellular components, highlighting plasma membrane– and vesicle-associated compartments.

In the dataset, we identified a total of 7001 proteins from Raji D1.3 B-EVs and 7148 from BAJB B-EVs, with high confidence (Fig. S2B; Supplementary Table SII). First, we compared our EV dataset with two curated datasets of EVs and secreted particles: Exocarta^42^ and Vesiclepedia^43^ (Fig. 3B). We found high overlap of our proteome and the curated datasets in both datasets but also identified additional 8272 proteins. The high number of non EV-linked proteins is likely to reflect our cell type, which is renowned for expressing a high amount of cell-type specific proteins^44–46^. Furthermore, the use of high-sensitivity data-independent acquisition proteomics enabled a much deeper coverage of the proteome allowing the capture of low-abundance proteins that may fall below the detection limits of curated studies in standard EV databases. Of note, we compared the data with a reference proteome of the species from which the surrogate antigen originated (donkey) to assess whether a potential contamination of the surrogate antigen itself persisted in the B-EV preparations. However, no antigen-derived peptides were detected in the dataset, suggesting that the EV purification protocol excludes possible antibodies and, thus, also secreted proteins below 150 kDa, from the culture medium (Supplementary Table SII).

We then performed an analysis for the differentially enriched (DE) proteins, to identify the proteins showing statistically significant (pooled p-value of ≤ 0.05) enrichment or reduction upon donor cell activation. B-EVs from 30-min antigen-activated Raji D1.3 cells showed 49 proteins with significantly increased (log2fold change >0.5), and 77 with significantly decreased levels (log2fold change <-0.5) (Fig 3C). BJAB EVs showed significantly increased levels for 11 proteins (log2fold change >0.5), and significantly decreased levels for 65 proteins.

In both cell lines, we also found several proteins exclusively either in B-EVs secreted from resting or activated cells (proteins present in ≥60% of EV replicates and in < 20% of WC replicates) (Fig. 3D, S2C), further supporting a shift in the type or content of B-EVs released upon cell activation. Interestingly, one of the proteins exclusively identified in B-EVs from activated Raji D1.3 is nuclear factor of activated T-cells, cytoplasmic 2 (NFATC2/NFAT1). NFATC2 has been shown to regulate the induction of gene transcription in lymphocytes during the immune response. The protein is known to be present in the cytosol and, upon TCR/BCR stimulation, to selectively translocate to the nucleus, where it forms transcription complexes with other components^47^. Based on our data, in addition to the nuclear translocation, NFAT2 would also get released via B-EVs upon antigenic stimulation. The presence of NFATC2 in Raji D1.3. B-EVs was also confirmed by western blot analysis (Fig. S2D).

We then performed KEGG pathway enrichment analysis on B-EVs from Raji D1.3 and BJAB, separately, to identify differentially represented biological pathways (Fig. 3E, F). B-EVs from resting Raji D1.3 showed an enrichment of proteins involved in glycosphingolipid biosynthesis, antigen processing and presentation pathways, and DNA replication, while the B-EVs released upon cell activation contained more proteins linked to RNA transport, ribosomes, and protein export (Fig. 3E, S2E). Regarding BJAB B-EVs, the analysis only revealed significant enrichment in the spliceosome pathway under resting conditions (Fig. S2F).

As antigen-induced activation of B cells triggers BCR signaling at the plasma membrane, we then asked if the BCR signaling proteins were also present in the B-EVs (Fig. 3G). While most BCR signaling proteins were found more abundant in donor cells, some proteins were enriched in the B-EVs from both cell lines (MAPK1, BLNK, PLCg2, BCAR1, PIK3CD, LPXN, BAX). Many of these proteins have been linked to actin cytoskeleton remodeling and calcium signaling. The differences identified in the B-EV content upon activation included more KLHL6 and PTPN22 in BJABs and more abundant BLNK, LCK, and exclusively NFATC2, in Raji D1.3 B-EVs. Finally, we performed an analysis of the cellular components (GO:CC) of proteins exclusively identified in Raji D1.3-derived EVs. The top enriched cellular components, highlighted the plasma membrane and vesicle-associated compartments, further supporting the plasma membrane origin of the B-EVs (Fig. 3H).

As EVs are known to also carry RNA cargo, which can influence the recipient cells, we next investigated the RNA content of the B-EVs. We now focused on Raji D1.3 B-EVs, which showed robust differences at the proteomic and lipidomic level upon antigen activation, and asked whether the RNA repertoire is also modulated by the donor cell activation.

Overall, Raji D1.3 B-EVs displayed a similar degree of heterogeneity in their RNA content as observed in the proteomic and lipidomic datasets, where the main source of variance was attributed to biological variability among EV preparations rather than to antigen stimulation (Fig. S2G). Enrichment analysis of consistently identified transcripts revealed functional categories related to DNA/RNA packaging and conformation changes, protein translation, lipid metabolism, and energy production (Fig. 4A–D). These pathways align with the concept of regulated formation of EVs that encapsulate proteins and nucleic acids in a regulated and signal-responsive manner.

**Figure 4.**
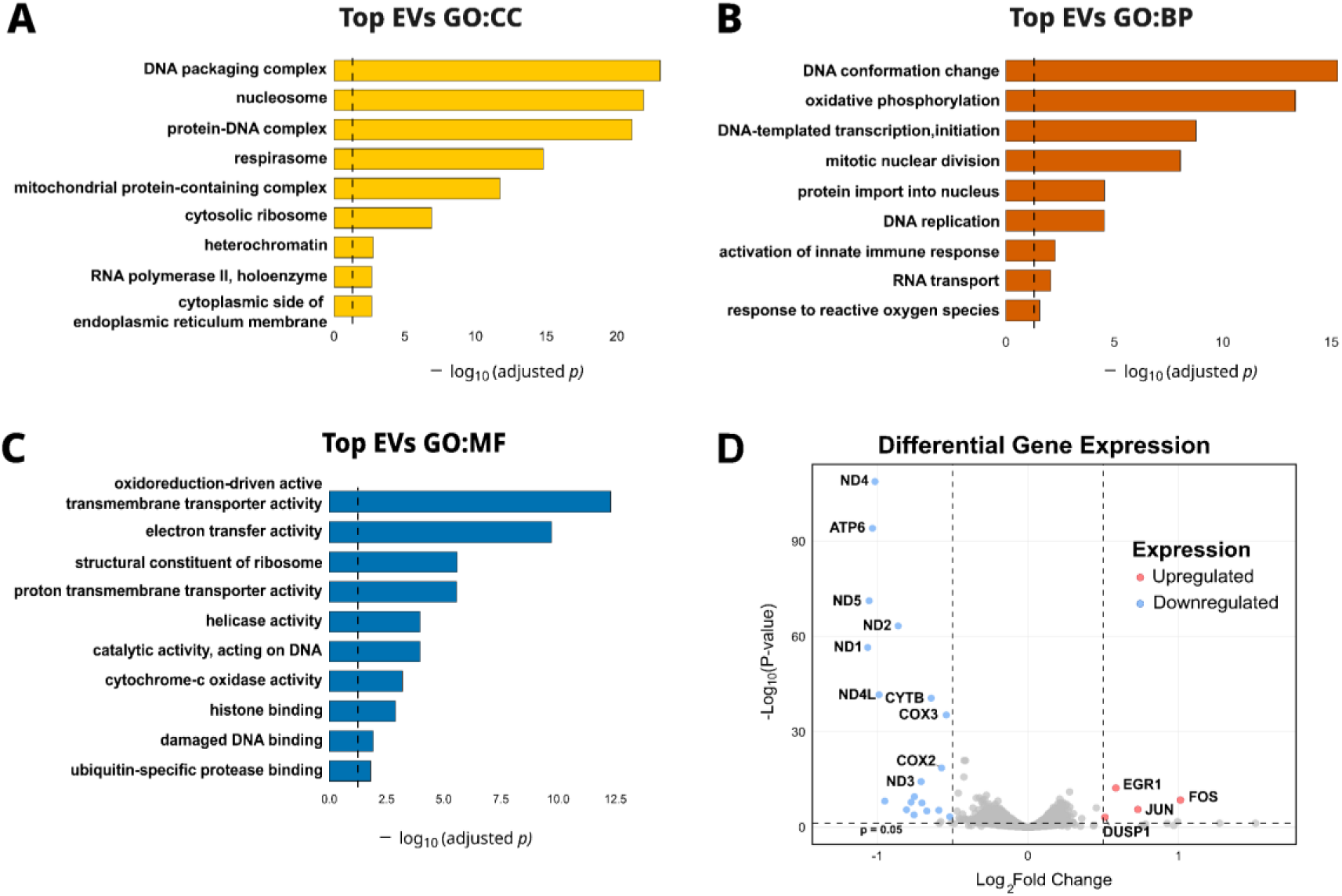
RNA content of the B-EVs is modestly altered upon antigenic activation. **(A-C)** Gene Ontology (GO) enrichment analysis of Raji D1.3 EV-associated transcripts. The genes corresponded to confidently identified RNA transcripts in isolated EVs from 30 minutes activated (ACT) or resting (non-activated, NA) cells, as earlier described. Bar plots show enriched terms (BH-corrected) in Cellular Components (CC) **(A)**, Biological Processes (BP) **(B)**, and Molecular Functions (MF) **(C)**. The bars represent -log10 (adjusted p-values) for selected representative terms. The dashed line indicates the significance threshold (adjusted p-value < 0.05). **(D)** Volcano plot of differentially expressed (DE) transcripts in Raji D1.3 EVs, isolated from cells that were activated or not. The plot displays log2 fold change (FC) (x-axis) versus -log10 p-value (y-axis) for ACT EVs genes compared to NA EVs genes. Colored points (right; red) represent significantly upregulated genes in ACT EVs (FC > 0.5, p < 0.05), colored points (left; blue) show significantly upregulated genes in NA EVs (FC < -0.5, p < 0.05), and grey points are non-significant. Dashed lines indicate significance thresholds. Top 10 most significantly up/down-regulated genes are labelled.

When specifically examining differentially represented transcripts between antigen-stimulated and control B-EVs, we detected a limited set of significantly up- or downregulated transcripts, suggesting that antigen activation, already at this timepoint of 30 minutes, induces subtle transcriptomic changes within the EV cargo. Nineteen transcripts were enriched in resting cell B-EVs while only 4 were enriched upon activation (Fig. 4D). This could indicate a selective inhibition in the delivery of certain transcripts into the budding EVs upon acute B cell activation, or that the diminution of these transcripts relates to shifts in the vesicle source. Most of the downregulated transcripts were mitochondrial, related to metabolism and energy production (ND4, ATP6, COX3)^48^, which could relate to the cells’ increased energy needs triggered by the BCR engagement and, thus, higher retention of these energy production -linked transcripts in cells. On the other hand, we identified only four upregulated transcripts in the B-EVs, namely FOS, JUN, EGR1, DUSP1. Interestingly, three of these four proteins, FOS, JUN, EGR1, are transcription factors.

Together, our multi-omics analyses demonstrated that B-EVs undergo modulation of their lipid, protein, and RNA composition upon acute antigenic activation of the donor cells.

### B-EVs interact with recipient cells

Differences in the composition of B-EVs shed by resting or antigen-activated cells suggest that they might also elicit distinct outcomes in recipient cells. First, we analyzed the interactions of B-EVs isolated as described above from the parental Raji D1.3 cell line. B-EVs were labelled with ExoGlow-RNA dye, and cell-associated fluorescence was monitored by confocal microscopy and flow cytometry at 4 and 20 h timepoints. Confocal microscopy revealed clear puncta of the RNA dye, indicating binding to or uptake of B-EVs by recipient cells (Fig. 5A). Flow cytometry analysis showed a clear increase in cell-associated RNA dye fluorescence in cells incubated with B-EVs compared with background signal from cells treated with the dye alone, supporting enhanced association of B-EVs with recipient cells. Cell-associated EV signal was detected already after four hours, and the proportion of EV+ cells approximately doubled following overnight incubation (Fig. 5B, S3A). B-EVs isolated from activated and resting cells were taken up at similar levels.

**Figure 5.**
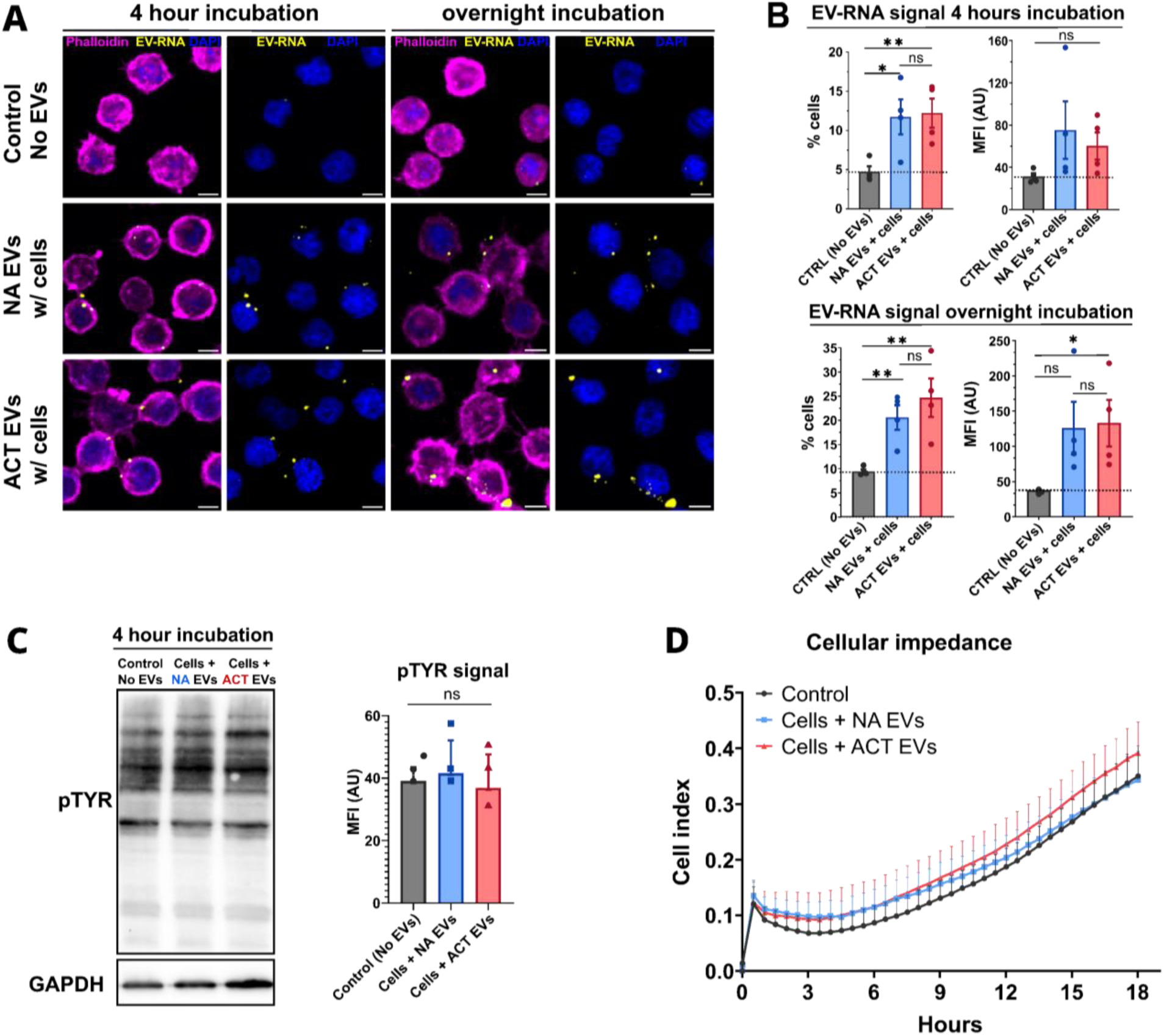
B-EVs interact with recipient B cells. **(A-B)** SEC-isolated EVs were stained with ExoGlow-RNA dye (5 uM) for 1 h, incubated with Raji D1.3 B cells for 4 hours or overnight, and imaged in 3D using a spinning disc confocal microscope (A). The representative images (sum of slices) depict dot-like EV-RNA signal associated with recipient cells; scale bar = 5 µm. (**B)** Quantification of ExoGlow-RNA dye signal was analyzed, by flow cytometry, in the recipient cells after incubation with B-EVs (4h, upper panel; overnight, lower panel). The graph on the left shows the number of ExoGlow-RNA dye positive cells, and the graph on the right shows the mean fluorescence intensity of the dye. Data is shown as mean ± SEM. Statistical significance was determined using two-tailed unpaired Student’s T test *p < 0.05, **p < 0.01, ***p < 0.001. **(C)** Western blot analysis and quantification of phospho-Tyrosine (pTyr) levels in recipient cells incubated with B-EVs from non-activated (NA) or 30-minute-activated cells for 4 hours. Data are presented as mean ± SEM from three independent experiments (N = 3). Statistical significance was determined using a two-tailed unpaired Student’s t-test. (**D)** Cellular impedance was measured in cells incubated with B-EVs from resting (non-activated (NA), or activated (ACT) cells for 18 hours (overnight). Cells were seeded onto fibronectin-coated E-View 96-well plates, followed by B-EV addition. Impedance was recorded every 30 minutes, and brightfield images were acquired every 45 minutes. (n=3).

We then investigated whether the B-EVs could influence the activation status of the recipient B cells. Total level of phospho-tyrosine, a common marker of BCR activation and B cell activation status, was assessed by Western blot, but no differences were detected between control cells and those incubated with B-EVs (Fig. 5C), suggesting that the uptake of B-EVs does not change the overall activity levels of the cells. To evaluate potential effects of B-EVs on recipient cell adhesion or morphology, we performed real-time cell impedance measurements that detect changes in electrical resistance arising as a sum of the cell number, adhesion, and morphology. B cells were plated on fibronectin-coated multi-well plates, mixed with B-EVs, and incubated for 18 hours. Impedance measurement and brightfield images were recorded every 30 and 45 minutes, respectively. As a sign of increased adhesion, cells treated with B-EVs from either activated or resting cells exhibited a slight increase in impedance relative to untreated control cells during the first 9 hours. In later time points, cells incubated with B-EVs from activated cells seemed to exhibit a moderate but consistent increase in their impedance (Fig. 5D). When we measured the recipient cell proliferation, we found no difference (Fig. S3B, C), suggesting that the increased impedance in cells receiving the activated B-EVs would result from higher adhesiveness or larger spreading morphology of the cells.

### B-EVs from antigen-activated B cells induce lytic reactivation of γ-herpesviruses

B cells are the natural site of latent infection for both human oncogenic γ-herpesviruses; EBV and KSHV^49^. These viruses are characterized by a biphasic life cycle with a silent, latent default program and an active, productive lytic replication which occurs sporadically following yet not completely understood triggers. Accumulating evidence suggests that B cell activation via the BCR pathway induces lytic reactivation of EBV and KSHV in latently infected cells^31,34^. Thus far, this has been hypothesized to be an autocrine effect, in which the virus is reactivated within the same cell in which the BCR pathway has been activated^31,34,50^. However, the hypothesis of a concomitant paracrine effect, in which secretions from activated B cells would stimulate lytic gene expression in infected neighboring cells, has, to our knowledge, never been tested. Given the differences observed in the composition of B-EVs from non-activated and activated B-cells, we asked whether the effect of B-EVs released from activated B cells could influence lytic reactivation of EBV and KSHV. To answer this question, we monitored the viral life cycle in the BJAB human B cell line, stably infected with the rKSHV.219 recombinant strain, for 72 hours after treatment with B-EVs isolated from either non-activated or activated Raji D1.3 cells.

The recombinant strain of KSHV, which harbors a constitutively expressed GFP gene and an RFP reporter that is expressed only during the lytic cycle, facilitating the fluorescent visualization of the viral latent-to-lytic switch^31,51^. Compared with PBS-treated cells, we observed an increase in RFP-positive cells upon B-EV treatment, which was even more pronounced when the cells were treated with the B-EVs isolated from activated cells (Fig. 6A, S4A). In treated cells, we also observed a clustered morphology, a common characteristic of lytic reactivation in infected lymphoblastoid cells^52,53^.

**Figure 6.**
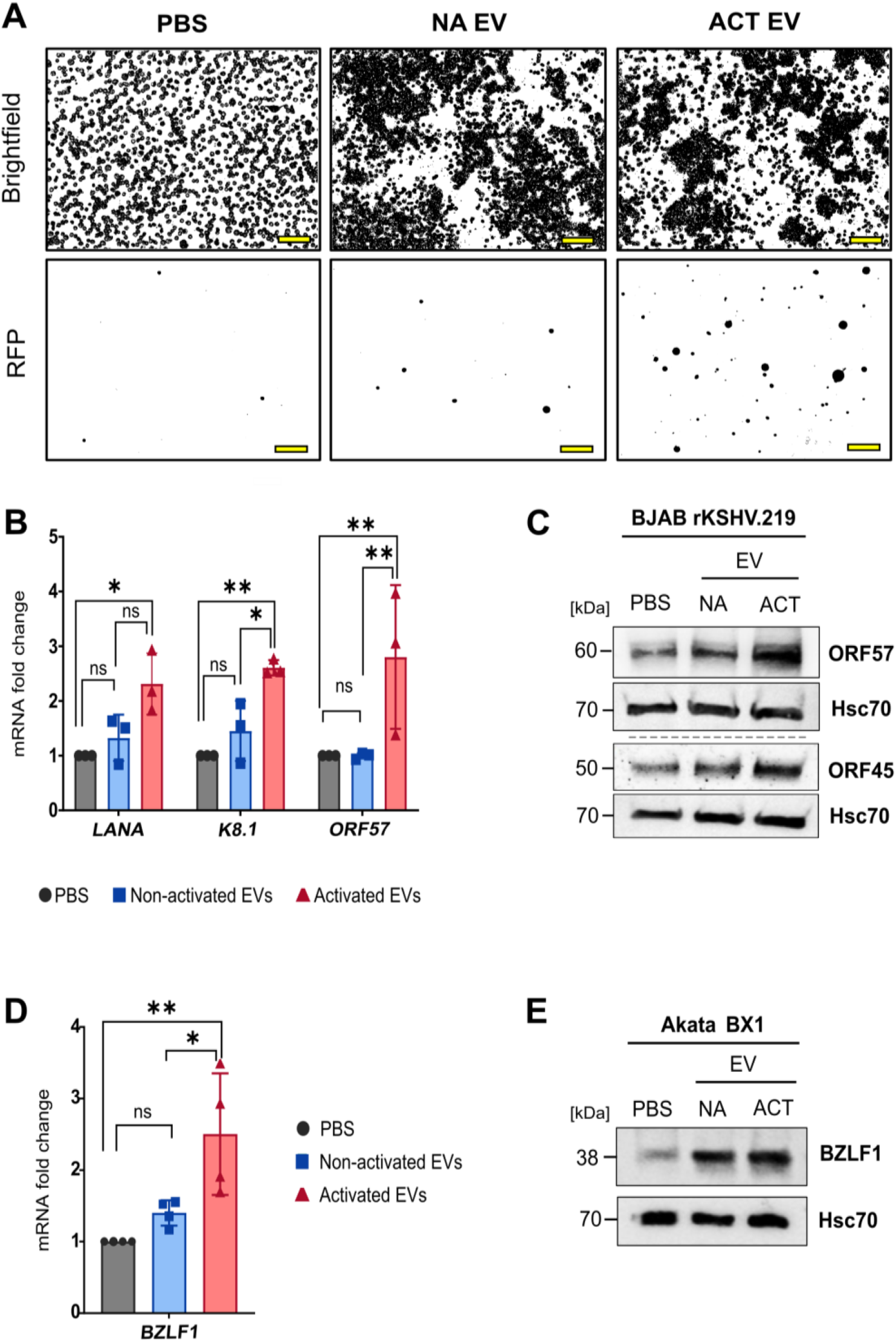
B-EVs originating from activated cells launch EBV and KSHV lytic gene expression. **(A-C)** BJAB rKSHV.219 cells were treated with 10 μg of B-EVs isolated from resting, nonactivated (NA) or 30 min activated (ACT) B cells for 72 hours. (**A)** Binary masks of Bright field (upper panels) and RFP fluorescent (lower panels) representative images (N=4). Scale bar = 100 μm. Binary masks applied with FIJI. (**B)** Transcript fold changes for the indicated viral markers. Actin was used as internal control and normalization value. Data are presented as mean ± SEM from three independent experiments (N = 3). Statistical significance was determined using two-way ANOVA and Tukey’s multiple comparisons correction **(C)** Representative immunoblot analysis of ORF57 and ORF45 protein levels. Hsc70 was used as loading control. (N=3). (**D, E)** EBV-positive AKATA cells were treated with 10 μg of EVs from NA B EV or ACT B EV for 72 hours. **(D)** BZLF1 transcript fold changes relative to PBS-treated control. 18S was used as internal control and normalization value. Data are presented as mean ± SEM from four independent experiments (N = 4). Statistical significance was determined using one-way ANOVA. **(E)** Representative immunoblot analysis of BZLF1 protein. Hsc70 was used as loading control. (N =3) *p < 0.05, **p < 0.01, ***p < 0.001.

To further characterize this effect, subsequent gene expression and immunoblotting analyses were performed to analyze the relative levels of viral lytic transcripts and proteins, respectively. To provide an overview of viral gene expression, we assessed levels of viral markers across the viral life cycle: latent (LANA), early lytic (ORF57 and ORF45), and late lytic (K8.1). The cells treated with B-EVs from activated cells exhibited a statistically significant increase in the expression levels of all viral genes tested when compared to the PBS control treatment, while the cells treated with B-EVs from resting cells showed no significant increase in viral gene expression (Fig. 6B). Immunoblot analysis confirmed the increase in ORF57 and ORF45 lytic protein levels upon treatment with B-EVs from activated cells (Fig. 6C, S4B). These results show that B-EVs secreted by activated B cells can induce significant changes in KSHV gene and protein expression, thereby facilitating lytic cycle activation in a paracrine manner.

Several reports have documented common molecular mechanisms of lytic reactivation shared by EBV and KSHV. The stress-related transcription factor heat shock factor 2 (HSF2), the mitochondrial protein TBRG4, and the cell cycle-related kinase PLK1 are recently identified examples of conserved regulators of both KSHV and EBV lytic cycles^54–56^. We next asked whether B-EVs would also stimulate EBV reactivation. To answer this question, we treated Akata BX1 cells, an EBV-infected human lymphoma B cell line, with B-EVs for 72 hours and analyzed EBV gene and protein expression. RT-qPCR analysis for the early lytic EBV gene BZLF1 revealed a significant increase in gene expression levels upon treatment with B-EVs from activated cells, while again, B-EVs originating from resting cells did not elicit significant changes (Fig. 6D). Immunoblot analysis showed upregulation of the product of BZLF1 gene, the protein Zebra1, already upon treatment with B-EVs from resting cells, but more pronounced in cells treated with B-EVs from activated cells (Fig. 6E, S4C).

Importantly, although we did not identify the activating surrogate antibody in the protein mass spectrometry analysis (see above), we still tested whether the lytic gene expression observed on the recipient cells could be due to possible residual surrogate antibody used to activate B-EV donor cells. For this, we treated the virally infected cells with either the surrogate antigen alone or with the surrogate antigen-containing cell-free culture medium that had been subjected to the EV isolation procedure (indicated as “Mock”). We observed that these treatments had no detectable effect, neither on the number of RFP positive cells nor on the KSHV ORF57 and EBV BZLF1 protein levels. This indicates that the reactivation of KSHV and EBV was not induced by leakage of the activatory antibody in our experimental systems (Fig. S4D-G). Furthermore, as Raji cells have been reported to contain latent EBV, we also analyzed our proteomic data against EBV reference sequences. We found EBV proteins in neither B-EV nor whole-cell samples, indicating that there is no leakage of viral proteins from the donor cells via B-EVs to the recipient cells.

Taken together, these results showed that B-EVs secreted by activated B cells facilitate lytic reactivation in KSHV and EBV positive B cells, thereby uncovering a novel stimulus for lytic reactivation in oncogenic human γ-herpesviruses and demonstrating that B cell activation can promote γ-herpesvirus lytic gene expression also in a paracrine fashion.

## Discussion

Here, we have performed a systematic characterization of B-EVs released from two human lymphoma-derived B cell lines, Raji D1.3 and BJAB. We show that, particularly B-EVs released from Raji D1.3 cells, are rapidly altered upon antigen signaling, leading to changes in their lipid, protein, and RNA content. Functionally, we found that B-EVs from antigen-activated cells trigger viral lytic gene expression in B cells harboring latent EBV or KSHV infections. These two human γ-herpesviruses have been described with oncogenic activity upon viral reactivation, yet, by largely unknown mechanisms. Our data suggest a novel paracrine mechanism for viral reactivation, by B-EVs secreted by nearby B cells that encountered their specific antigens.

While B cells have been reported to actively secrete EVs, the characterization of their content and functional properties has remained limited^16^. The effects of antigen-induced BCR signaling on EV release, composition, origin, and function, have not been previously addressed. We observed a clear modulation of the B-EVs secreted from RajiD1.3 cells already after 30 min of antigen activation, as evidenced by changes in the lipidome and proteome profiles as well as in the mRNA content of the vesicles. BJAB-derived B-EVs also showed modulation upon antigenic stimulus, but to a lesser extent. The lipidomic profile of our B-EVs suggests that B cells primarily secrete EVs that bud from the plasma membrane (Fig. 2). To our knowledge, this is the first lipidomic analysis of B-EVs, establishing a reference for future studies of B-EVs from different B-cell types. Treatment of donor B cells with CD24 agonist, known for its apoptosis-inducing and proliferation-suppressive effects, has been shown to increase the membrane protein content in B-EVs and elicit enhanced signaling responses in the recipient B cells^20,57,57,58^. Ayre et al (2015)^59^ propose that B cells could respond to the CD24 engagement already in 15 min, however, the analyses of the EVs remained somewhat ambiguous. Subsequently, the same lab performed a more thorough B-EV analysis after 1 h of anti-CD24 stimulation. These publications support our findings that B cells are primed to respond to signals with immediate modulation of B-EV secretion. Similarly, the work from Christian lab proposing that B cells primarily respond by increasing the release of plasma membrane-derived ectosomes^58^, aligns well with our lipidomic profiling, proposing an enhancement in the plasma membrane budded vesicle content upon cell activation. While some proteinaceous material was also detected in our preparations, often bound to the B-EV surface, the contribution of non-EV material in our analyses cannot be excluded. However, the differential lipid, protein, and RNA content, together with the functional effects of B-EVs from antigen-activated cells, indicate that vesicles are the primary mediators of these effects.

EBV and KSHV establish lifelong latent infections in B cells. Upon appropriate stimuli, a subset of these latent cells can reactivate and initiate lytic gene expression. The switch to the lytic cycle is driven by the immediate-early genes EBV BZLF1 and KSHV RTA, which act as master regulators of the downstream lytic program. In our study, we detected an increase in lytic gene expression following treatment with B-EVs from activated cells (Fig. 6) suggesting that differentially expressed components present in these B-EVs contribute to viral reactivation. A conserved molecular feature of this process is the involvement of cellular transcription factors that respond to stress and signaling cues, such as the activator protein 1 (AP-1). AP-1 is a transcription factor complex composed of heterodimers of mainly c-Jun and Fos family proteins, which belong to the immediate early genes, and are commonly upregulated in response to various growth factor and stress stimuli. Notably, our transcriptomic profiling revealed a significant upregulation of both c-Jun and Fos in B-EVs from antigen-stimulated cells (Fig. 4), and FosB, a homologue of Fos, protein was exclusively identified in B-EVs from activated cells (Fig. 3). Both KSHV RTA and EBV BZLF1 promoters contain AP-1 consensus sequences, and it has been demonstrated that the recruitment of AP-1 complex to these viral promoters is required for the initiation of the lytic cycle^60,61^. This suggests that increased expression levels of these proteins in recipient cells may contribute to the effect of B-EVs on viral reactivation.

The possibility of paracrine regulation of the viral life cycle by neighboring antigen-activated B cells suggests that an inflammatory environment could be sufficient to stimulate viral activation in bystander cells with unrelated antigen specificities. Given that more frequent lytic reactivation has been identified as a risk factor for the development of γ−herpesviruses-associated malignancies, our findings suggest a molecular basis, by which virus-infected patients with history of chronic local inflammatory symptoms are exposed to a greater risk of developing γ−herpesviruses-associated tumours^31,50^. For instance, Kaposi’s sarcoma, an angiogenic tumor of the skin, caused by KSHV occurs often at sites of previous trauma, including scars and inflammatory lesions^62,63^. Additionally, one of the risk factors for Epstein-Barr virus (EBV)-associated gastric cancer (EBVaGC) is recurring gastritis, indicating that a local inflammatory environment facilitates the insurgence of this type of cancer. Another interesting epidemiological evidence is that remnant gastric cancer, i.e. a rare gastric cancer that occurs in sites of previous gastrectomy shows a significantly higher association with EBV compared to other types of gastric cancer, again indicating that herpesvirus-driven cancers occur more frequently in the sites of previous damages^64^.

Our results demonstrate increased lytic gene expression after treatment with B-EVs from activated B cells. This highlights a novel paracrine route for triggering the latent-to-lytic switch, as lytic reactivation following BCR activation was previously thought to be an autocrine process^31^. Further investigation on the molecular mechanisms involved in the paracrine regulation of the γ-herpesvirus life cycle could advance our understanding of viral tumor microenvironments and potentially have implications in the development of novel EV-based therapies.

## Materials and methods

### Cells

Raji D1.3 (a subclone of Raji D1.3, a Burkitt’s lymphoma cell line) expressing the transgenic IgM B-cell receptor D1.3 specific to hen egg lysozyme, and BJAB (Burkitt’s lymphoma cell line) obtained from DSMZ (German Collection of Microorganisms and Cell Cultures), were maintained under specific conditions. Raji D1.3 D1.3 cells were maintained at 37°C and 5% CO_2_ in complete RPMI-1640 (Cytiva) supplemented with 10% fetal calf serum (FCS)(Biovest), 1% L-glutamine (Euroclone), and 1% Penicillin-Streptomycin (PenStrep) (Euroclone). BJAB cells were cultured in cRPMI supplemented with 20% FCS, 1% L-glutamine, and 1% PenStrep.

The virus-infected cell lines used were BJAB rKSHV.219 and Akata BX1, both of which originate from Burkitt’s lymphoma patients^33,65^ and were subsequently infected with recombinant viral strains in culture. The BJAB rKSHV.219 cell line is stably infected with a strain of KSHV called rKSHV.219, and was kindly provided by Thomas F. Schultz at Hannover Medical School^31^. The rKSHV.219 strain has been genetically modified to harbor a puromycin resistance gene, an RTA-inducible RFP and a constitutively expressed GFP. These characteristics allow antibiotic selection of cells to maintain viral presence in the cells, as well as viral life cycle monitoring through fluorescence^66^. The Akata BX1 cell line is infected with a recombinant strain of EBV called BX1, which harbors a puromycin resistance gene and a constitutively expressed GFP (Molesworth et al., 2000). The cell line was kindly provided by Rob White at Imperial College London. These cell lines were maintained at 37ᵒC and 5% CO2, using cRPMI-1640 (Sigma-Aldrich) supplemented with 10% heat-inactivated fetal bovine serum (Serana), 1% L-glutamine (Sigma-Aldrich) and 1% Penicillin-Streptomycin (Sigma-Aldrich).

### Isolation of extracellular vesicles (EVs)

Extracellular vesicles from Raji D1.3 and BJAB cells were isolated using size exclusion chromatography (SEC) with qEV original 35 columns (Izon). Initially, cells were collected by centrifugation (1400 rpm, 3 min), washed twice with serum-free RPMI medium, and resuspended in 2 ml of serum-free RPMI. Half of the cells were activated with surrogate antigen (10 µg/mL of donkey anti-mouse IgM-Fab^2^ for Raji D1.3 and goat anti-human IgM-Fab^2^ for BJABs) while half was left resting. After a 30-minute incubation period, supernatants were collected after centrifugation steps at 1500 g and 10,000 g for 10 min at +4°C. The supernatant was loaded onto the SEC column and EVs were eluted with filtered PBS as per the manufacturer’s protocol. The first 4 mL of eluate were discarded, and the subsequent 10 mL of EV-enriched content were pooled for protein concentration using 15 mL Amicon protein concentrator tubes 30 kDa MWCO (Sigma Aldrich), supplemented, if necessary, with an additional 500 µl in Amicon protein concentrator PES with a 3K MWCO filter (Sigma Aldrich). Protein concentration was determined using a BCA assay, and samples were used for characterization and functional assays. The samples that were subjected to lipidomic analysis, were not concentrated with the Amicon filters and were instead subjected to ultracentrifugation (described later).

### Protein measurements

Following the manufacturer’s instructions, protein concentrations in isolated extracellular vesicles and whole-cell lysate samples were determined using the Pierce 660 nm Protein Assay (Thermo Fisher Scientific) in combination with the Ionic Detergent Compatibility Reagent (Thermo Fisher Scientific).

### Nanotracking analysis (NTA)

The size distribution and concentration of SEC-isolated EVs were measured using the ZetaView PMX-120 system (Particle Metrix) at the EV Core Facility of the University of Helsinki (FIMM EV Core), following their standard operating procedures. Measurements were conducted at 22 °C across 11 positions, using the following settings: minimum area = 10, maximum area = 1000, and minimum brightness = 30. A total of four biological replicates were analyzed. Raw NTA data was processed using Microsoft Excel. Data smoothing was performed using a rolling average, and the resulting curves were visualized as scatter plots in Microsoft Excel.

### Transmission electron microscopy

For electron microscopy, 5 µL of EV samples in PBS were incubated on hydrophilized formvar/carbon-coated copper mesh grids for 1 minute. After removal of excess sample, grids were washed thrice with an equal volume of milliQ water and negative-stained with 5 µL of 2% uranyl acetate for 1 minute. Post-staining, excess solution was blotted off and the grids were left to dry at room temperature for 10 minutes. Sample grids were stored in the grid box and imaged within a week of preparation using the JEM-1400 Plus transmission electron microscope equipped with OSIS Quemesa 11 Mpix bottom mounted digital camera.

### Immunoblotting

Western blot analysis was performed on EVs and whole cell lysates (WCs) from non-activated (NA) and activated (ACT) Raji D1.3 and BJAB cells using either 4-15% gradient or 10% precast polyacrylamide gels under non-reducing conditions. WCLs were prepared in Triton X-100 lysis buffer (1% Triton X-100, 50 mM Tris-HCl (pH 7.4), 150 mM NaCl, 1 mM EDTA, and protease/phosphatase inhibitors), sonicated (30 sec on/30 sec off cycles, high frequency), and boiled for 8 min at 95–97 °C in 4× SDS Laemmli buffer without β-mercaptoethanol, same as for EV samples. Equal protein amounts (∼5 µg) were loaded, run, and transferred to nitrocellulose membranes using Trans-Blot Turbo Mini Transfer Packs (Bio-Rad). Membranes were blocked with 5% milk in 1xTBS-Tween 0.05% (TBS-T) for 1 h, incubated overnight at 4 °C with primary antibodies (see table), washed in TBS-T, incubated with HRP-conjugated secondary antibodies in 5% milk in TBT-T for 1h at RT and developed using ECL substrate (1:1 ratio). Signal was detected with the ChemiDoc MP Imaging System (Bio-Rad).

For viral reactivation studies; cell pellets were lysed in 3× Laemmli buffer (30% glycerol, 187.5 mM SDS, 3% Tris-HCl, 0.015% bromophenol blue, 3% β-mercaptoethanol) and boiled at 95 °C for 10 min. Lysates were resolved on 4–20% Mini-PROTEAN TGX Stain-Free Gels (BIO-RAD) alongside Precision Plus Protein Dual Colour Standards (BIO-RAD), and transferred to Amersham Protran 0.45 μm nitrocellulose membranes (Cytiva) using the Trans-Blot Turbo system (BIO-RAD). Membranes were blocked in 5% milk in PBST (0.1% Tween-20 in PBS) for 1 h at RT, washed, and incubated overnight at 4 °C with primary antibodies (see table) diluted in 0.5% BSA, 0.02% NaN₃ in PBS. After washing, membranes were incubated with secondary antibodies in 5% milk/PBST for 1 h at RT, washed again, and developed using SuperSignal West Pico PLUS substrate (Thermo Fisher). Signal was detected with the iBrightCL1000 system (Thermo Fisher), and Hsc70 was used as a loading control. Band intensities were quantified using Fiji ImageJ.

### Quantitative mass spectrometry-based shotgun lipidomics

For lipidomic analysis of the B-EV from RajiD1.3 and BJAB cells, 8,5 ml out of total 10 ml of the pooled supernatants (in PBS, as described earlier) were collected to polypropylene tubes (331372; Beckman Coulter) and subjected to ultracentrifugation (Beckman Coulter) at 100000 g (90’, +4 Celsius) using swinging-bucket rotor SW41. The supernatant was carefully decanted and removed with pipette and the tubes containing thin pelleted samples were stored at -80 Celsius until lipidomic analysis. For whole cell lipidomic analysis of the same cells from where EVs were collected, one million cells from all conditions were collected into separate tubes, centrifuged (1300 rpm, 3’), cells washed with Milli-Q, centrifuged (1300 rpm, 3’) and cell pellets stored in -80 C until lipidomic analysis. Quantitative mass spectrometry-based shotgun lipidomics was performed as previously described in Nielsen et al ^37^, with minor modifications. The lipid extraction was carried out at 4°C using HPLC-grade solvents in Eppendorf tubes. Raji D1.3 D1.3 and BJAB cells (∼1 million cells) and the corresponding B-EV samples collected by ultracentrifugation were mixed with 155 mM ammonium bicarbonate (AB, Sigma-Aldrich, FL40867) to a final volume of 200 µl. Each sample was spiked with an internal lipid standard mixture. Chloroform/methanol (C/M, 2:1, v/v; 1 ml) was added, and samples were shaken for 20 min (2,000 rpm) before centrifugation (1 min, 500 g). The lower organic phase was transferred to a new Eppendorf tube, and evaporated for at least 75 min using a vacuum concentrator. The resultant lipid film was dissolved in 100 µl C/M 1:2 (v/v) by shaking (5 min, 2,000 rpm) and centrifuged (15 min, 18,000 g) to remove insoluble particles. Extracted lipids were then mixed with positive and negative ionization solvents (13.3 mM AB in 2-propanol or 0.2% (v/v) methylamine in methanol, respectively) and analyzed in the positive and negative ion modes on Orbitrap Fusion mass spectrometer (Thermo Fisher Scientific) equipped with TriVersa NanoMate (Advion Biosciences) for automated and direct nanoelectrospray infusion. Lipid identification and extraction of precursor- and fragment-ion intensities were carried out using LipidXplorer version 1.2.4^67^, and lipid quantities in samples were calculated using the R-based in-house software LipidQ (https://github.com/ELELAB/lipidQ). Statistical analyses were carried out in GraphPad Prism 10.4.2, and heatmaps were produced using Morpheus, https://software.broadinstitute.org/morpheus.

### Protein mass spectrometry

#### Sample preparation and protein digestion for LC/MS-MS

Samples containing 85 μg of total EV or 70 μg of total WC proteins were further processed by the filter-aided sample preparation (FASP) method^68^. In short, EVs isolated from Raji D1.3 and BJAB or WCLs were lysed in 5x SDS-lysis buffer [2% (w/v) SDS, 500 mM Tris/HCL pH 7.6, 0.5 M DTT]. For each sample, the appropriate amount of lysate was mixed with UA buffer (8 M urea in 0.1 M Tris-HCl, pH 8.5) in Microcon YM-10 centrifugal filter units (Merck Millipore) and centrifuged at 14,000 × g at room temperature (RT) for 40min. The retained proteins were washed once with UA buffer and centrifuged again at 14,000 × g at RT for 40min. Proteins were then alkylated with 0.05 M iodoacetamide (Sigma-Aldrich) freshly dissolved in UA buffer, and incubated in the dark for 30 minutes at room temperature. Samples were washed twice with UA and twice with DB buffer (1 M urea in 0.1 M Tris-HCl, pH 8.5) before transferring the filter to a fresh collection tube.

Proteins were digested on-filter using Lys-C/Trypsin mix (Promega) at an enzyme-to-protein ratio of 1:25. Digestion was carried out for 18h at 37 °C. Peptides were eluted by sequential centrifugation with DB buffer, acidified with formic acid (Thermo Fisher Scientific) to a final concentration of 1% (final pH 2–3), and clarified by centrifugation at maximum speed for 15 minutes. Peptides were desalted using Sep-Pak tC18 well plate (Waters). Dried peptide samples were dissolved into 0.1% formic acid and peptide concentration was measured using Qubit Flex (Thermo Fisher Scientific). Based on concentration measurement, 1 µg of each sample was injected for analysis. Washes were submitted between each sample to reduce potential carry-over of peptides.

#### LC-ESI-MS/MS Analysis

The LC-ESI-MS/MS analyses were performed on a nanoflow HPLC system (Easy-nLC1000, Thermo Fisher Scientific) coupled to the Orbitrap Exploris 480 mass spectrometer (Thermo Fisher Scientific, Bremen, Germany) equipped with a nano-electrospray ionization source and FAIMS interface. Compensation voltages of -40 V and -60 V were used. Peptides were first loaded on a trapping column and subsequently separated inline on a 15 cm C18 column (75 μm x 15 cm, ReproSil-Pur 3 μm 120 Å C18-AQ, Dr. Maisch HPLC GmbH, Ammerbuch-Entringen, Germany). The mobile phase consisted of water with 0.1% formic acid (solvent A) or acetonitrile/water (80:20 (v/v)) with 0.1% formic acid (solvent B). A 120 min step gradient (from 5 to 21% of solvent B in 62 mins, from 21% to 36% of solvent B in 48 mins, from 36% to 100% of solvent B in 5 min, followed by a 5 min wash stage with 100% of solvent B) was used to eluate peptides. Samples were analysed by a data independent acquisition (DIA) LC-MS/MS method. MS data was acquired automatically by using Thermo Xcalibur 4.6 software (Thermo Fisher Scientific). In a DIA method a duty cycle contained one full scan (400–1000 m/z) and 30 DIA MS/MS scans covering the mass range 400–1000 with variable width isolation windows.

### Protein Identification and quantification

Raw data files were analyzed using Spectronaut (v19.9, Biognosys AG) with the DirectDIA approach was used to identify proteins and label-free quantifications were performed with MaxLFQ. Protein identification was performed using Pulsar search with a Trypsin/P enzyme specificity, allowing up to 2 missed cleavages. Carbamidomethylation was set as a fixed modification, while oxidation (M) and Acetyl (protein N-term) were considered variable modifications. Database search was done against a SwissProt Homo sapiens reference proteome (Swiss-Prot 2025_01 Homo Sapiens) supplemented with common contaminants provided by the Biognosys default contaminant database, a donkey proteome reference (UniprotKb 2026_01 Equus asinus) and two EBV proteome reference (Swiss-Prot2026_01 Epstein-Barr virus (strain B95-8) (HHV-4) (Human herpesvirus 4) and Swiss-Prot 2026_01 Epstein-Barr virus (strain AG876) (HHV-4) (Human herpesvirus 4)). Peptide-spectrum match (PSM), peptide, and protein group false discovery rates (FDRs) were all controlled at 1%, using a decoy-based approach with mutated decoys. Protein quantification was performed based on MS2-level extracted ion chromatogram (XIC) and normalized based on RT dependent local regression model described by Callister et al^69^.

Identified proteins from Spectronaut (Biognosys) imported into in R statistical programming language for post-processing and statistical analysis. Post-processing steps included the removal of contaminants (common contaminants, proteins with lower than 2 unique peptides and a Q-value >0.01) to ensure high-confidence identifications. Separate datasets were generated for BJAB and Raji D1.3 cell lines comparing EV and WCL fractions. Identified protein proteins were then classified as either exclusive or differentially expressed (DE) based on the number of valid values (non-missing values) per condition. Proteins detected in only one condition were considered exclusive. In contrast, proteins with ≥ 60% valid values for every condition were retained for differential expression analysis. Missing values were imputed using k-nearest neighbor (k-NN) imputation (k = 10) from the impute R package. For Log₂ fold change values were calculated for each protein by averaging the difference in signal intensity between five biological replicates of activated vs. non-activated EVs. A student’s T-test was applied to the replicate-level log₂ differences to assess statistical significance under the null hypothesis that mean difference equals zero.

### EV RNA extraction and library preparation/sequencing

Total RNA was extracted from freshly isolated EVs using the exoRNeasy Midi Kit (QIAGEN) according to the manufacturer’s instructions. RNA concentration was measured with the Qubit RNA BR Assay Kit and Qubit 4 Fluorometer. RNA samples (100 ng/µL) from three biological replicates of EVs derived from non-activated and activated cells were then processed by the Finnish Functional Genomics Centre (Turku Bioscience). Library preparation was performed using the Illumina Stranded Total RNA Library Preparation protocol (Illumina, 1000000124514) and the Illumina Stranded Total RNA Preparation Ligation Kit (Illumina). Sample quality was assessed with an Agilent Bioanalyzer and RNA concentration was determined using the Qubit®/Quant-IT® Fluorometric Quantification Kit (KAPA Biosystems). Sequencing was carried out with the NovaSeq 6000 platform SP v1.5 flow cell, 2 × 100 bp read length, one lane/pool, and a 1% PhiX v3 control spike-in. The typical data yield was 650–800 M reads per run (two lanes) from the NovaSeq 6000 SP run. Raw sequencing data were converted to fastq format using the bcl2fastq2 conversion software (Illumina).

### RNA transcriptomics

Data cleaning, filtering, and analysis were performed in R. Raw gene counts were processed with the DESeq2 package for differential expression analysis. Genes with fewer than 100 total counts across all samples were excluded. Sample metadata including experimental conditions and biological replicates was used to define the experimental design. Normalization was performed using DESeq2’s median-of-ratios method, which estimates size factors to account for library size differences. Quality control included assessments of library size, gene expression dispersion, and size factor distribution.

Pairwise differential expression analysis was performed using the Wald test, with contrasts defined by the experimental design. LFCs were shrunk using the shrinkage estimator function in DESeq2 to mitigate the influence of genes with low counts or high dispersion, without affecting the adjusted p-values (FDR).

Plots were generated with ggplot2, cowplot, and pheatmap packages.

### Functional enrichment analysis (proteomics and transcriptomics)

Principal Component Analysis (PCA) was performed using scaled log₂ intensity values, with missing values imputed as zeros from FactoMineR package. PCA plots are shown with samples colored by condition and 95% confidence ellipses shown. Heatmap was constructed using hierarchical clustering (Ward.D2 method, Euclidean distance), grouping proteins into 10 clusters. PCA plots and Heatmaps were generated using the package ggplot2.Over-representation analysis (ORA) of Gene Ontology (GO) terms and KEGG pathways was performed using significant proteins/genes from either differential expression (p < 0.05) or exclusive detection criteria. ORA was conducted using the clusterProfiler package (v4.10.0) with UniProt identifiers, querying the org.Hs.eg.db database. Terms were considered significant at adjusted p < 0.05 (Benjamini-Hochberg). For KEGG pathway analysis, significantly up- and downregulated proteins (based on log₂ fold change and p < 0.05) were separated to visualize top EV pathways upon activation. Venn diagrams were generated using the ggVennDiagram and eulerr packages.

### Microscopy and image acquisition

Following incubation with RNA-stained EVs, cells (30 000/well) were resuspended in imaging buffer (0.5% FCS in PBS), plated onto fibronectin-coated (4 µg/mL) 12-well microscopy slides and incubated for 30 minutes at 37 °C in 5% CO₂. Cells were then fixed in 4% paraformaldehyde (PFA) for 10 minutes at room temperature (RT), washed with PBS, and stained with phalloidin-AF555 (1:200 dilution) in 1% BSA/PBS containing 0.3% Triton X-100 for 1 hour at RT. After staining, samples were washed three times with PBS and mounted using Fluoromount with DAPI. Slides were left to dry at RT, protected from light overnight. Image acquisition was performed using a 3i Marianas spinning disk confocal microscope (CSU-W1, 50 µm pinholes; Yokogawa) controlled via SlideBook 6 software. A 63× Zeiss Plan-Apochromat oil immersion objective (NA = 1.4; working distance = 0.19 mm) was used, and images were captured with a Hamamatsu sCMOS Orca Flash4.0 camera (2048 × 2048 pixels, 6.5 × 6.5 µm). Excitation was achieved using lasers at 405 nm (100 mW), 488 nm (150 mW), and 561 nm (100 mW). For 3D z-stack imaging, optical sections were collected at 300 nm intervals. Image processing and analysis were conducted with FIJI (ImageJ).

### Flow cytometry

Cells incubated with EVs were collected by centrifugation at 1400 rpm for 3 minutes at 4 °C and fixed in 4% paraformaldehyde (PFA) for 10 minutes at room temperature (RT). After fixation, samples were transferred to a V-bottom 96-well plate and centrifuged at 1500 rpm for 2 minutes at 4 °C. Permeabilization, blocking, and staining were performed in 1% BSA with 0.1% Triton X-100 in PBS. Cells were incubated with primary antibodies for 1 hour at RT. Following primary antibody incubation, samples were washed with PBS, centrifuged, and incubated with secondary antibodies for 1 hour at RT in 1%BSA/PBS. After staining, cells were washed twice with PBS and resuspended in 1% BSA/PBS. Samples were stored at 4–8 °C until data acquisition. Flow cytometry was performed using a BD LSRFortessa, and data was analysed with BD FACSDiva Software v8 and FlowJo v10. Statistical analysis and graphs were done in GraphPad Prism 8.

### Impendence measurement

To assess the effect of EVs on recipient cell behavior and adhesion in real time, we used the xCELLigence RTCA eSight system (Agilent) with RTCA E-Plate View 96-well plates. Wells were pre-coated with fibronectin (6 µg/mL) for 2 hours at 37 °C, washed twice with PBS, and kept in PBS until use. Raji D1.3 cells (50,000 cells/well) were seeded in the presence of 1 µg of EVs or w/o EVs (Control) and incubated for 18 hours at 37°C in 5% CO^2^. Cellular impedance was measured every 30 minutes, and brightfield images were captured every 45 minutes throughout the incubation. Raw Cell Index (CI) values were exported and analyzed in Microsoft Excel. Average values were calculated from technical duplicates, and impedance curves were plotted using GraphPad Prism.

### Incubation of EVs with cells

For monitoring of EVs interaction with cells, 2 × 10⁶ of Raji D1.3 cells were incubated with 40 µg of EVs in a 6-well plate in cRPMI. Cells were given either EVs from non-activated cells (NA EVs), 30-minute-activated cells (ACT EVs), or PBS alone (control, no EVs). Incubations were performed for 4 hours or overnight under standard culture conditions. When using smaller well formats or different experiment scales, cell and EV quantities were proportionally adjusted to ensure consistency and reproducibility. Prior to EV interaction assays, approximately 85 µg each of NA and ACT EVs were labelled using the ExoGlow™-RNA EV Labeling Kit (System Biosciences), following the manufacturer’s protocol. Briefly, EVs were mixed with 5 µM of RNA-specific fluorescent probe and an equal volume of incubation buffer (1:1 ratio). The mixture was incubated for 1 hour at 37 °C in the dark. Labelled EVs were then added to cell cultures and incubated for 4 hours or overnight. Control cells were incubated only with RNA probe mixture without EVs. Cells were then collected and processed for flow cytometry and confocal microscopy.

To analyze the effect of EVs from activated vs. non-activated cells on virus-infected cells, BJAB rKSHV.219 and Akata BX1 cells were seeded in 24-well-plates in a density of 1 x 10^6^ cells/ml. Cells were treated with 10 μg/well of EVs from either non-activated B-cells (NA EVs) or from activated B-cells (ACT EVs). Control cells were treated with PBS alone, equivalent to the given volume of EV-solution. As additional controls, cells were either treated with the surrogate antigen used for B-cell activation (10 µg/mL of donkey anti-mouse IgM-Fab2), or with the surrogate antigen complemented cell-free culture medium that had been subjected to the EV isolation procedure (indicated as “Mock”). Cells were then incubated at 37ᵒC and 5% CO2 for 72 hours. Before harvesting, the cells were monitored with a ZOE Fluorescent Cell Imager (Bio-Rad) instrument, using the brightfield, GFP and RFP channels. Upon harvesting, the cells were stored as pellets at -80ᵒC and used for subsequent analysis.

### RT-qPCR

RNA was extracted from frozen cell pellets using the RNeasy Plus Mini Kit (QIAGEN), and the yield was measured by NanoDrop 2000 spectrophotometer (Thermo Fisher Scientific). 900 ng of RNA per sample was reverse-transcribed to synthesize cDNA, using the iScript Reverse Transcription Supermix (Bio-Rad) and incubated in a thermocycler according to the manufacturer’s instructions. The RT-qPCR reactions were carried out using SensiFAST SYBR Hi-ROX kit (Meridian Bioline) in QuantStudio 3 Real-Time PCR systems (Applied Biosystems, Thermo Fisher Scientific). Samples were run in triplicates, and assuming 100% efficiency of cDNA synthesis, final RNA concentration was 18 ng/well. Actin or 18S was used for internal controls and normalization. At least three biological repeats were performed for each experiment. Primer sequences used are listed in the table below.

**Table 1.**
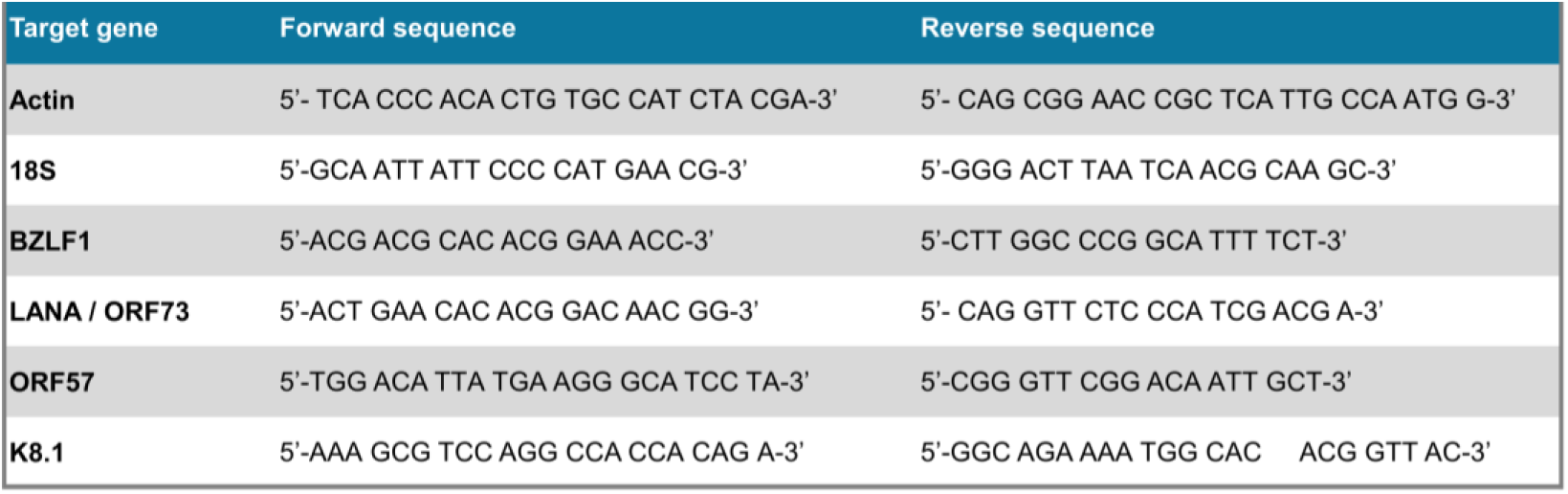
Primer sequences used for RT-qPRC.

### Statistical analyses

Data generated using the methods above were statistically analyzed using GraphPad Prism 8 and 10.4.2. Unless differently stated in the figure legends, each graph shows individual values of biological replicates as symbols and the mean as bars. Error bars indicate either standard deviation (SD) or standard error of the mean (SEM) as specified in the individual figure legends. For immunoblot quantifications and flow cytometry analysis statistical significance was determined using two-tailed paired/unpaired Student’s T test *p < 0.05, **p < 0.01, ***p < 0.001. For viral lytic gene reactivation experiments, Analysis of variance (ANOVA) tests were performed on all datasets, either as ordinary one-way ANOVA (1 variable per condition) or two-way ANOVA (≥2 variables per condition). Additionally, Tukey’s multiple comparisons test was performed to measure the effects of treatments and conditions between samples. Alpha limits were set as 0.05, and depicted as follows: *: p < 0.05; **: p < 0.01; ***: p < 0.001; ns: non-significant.

**Table 2.**
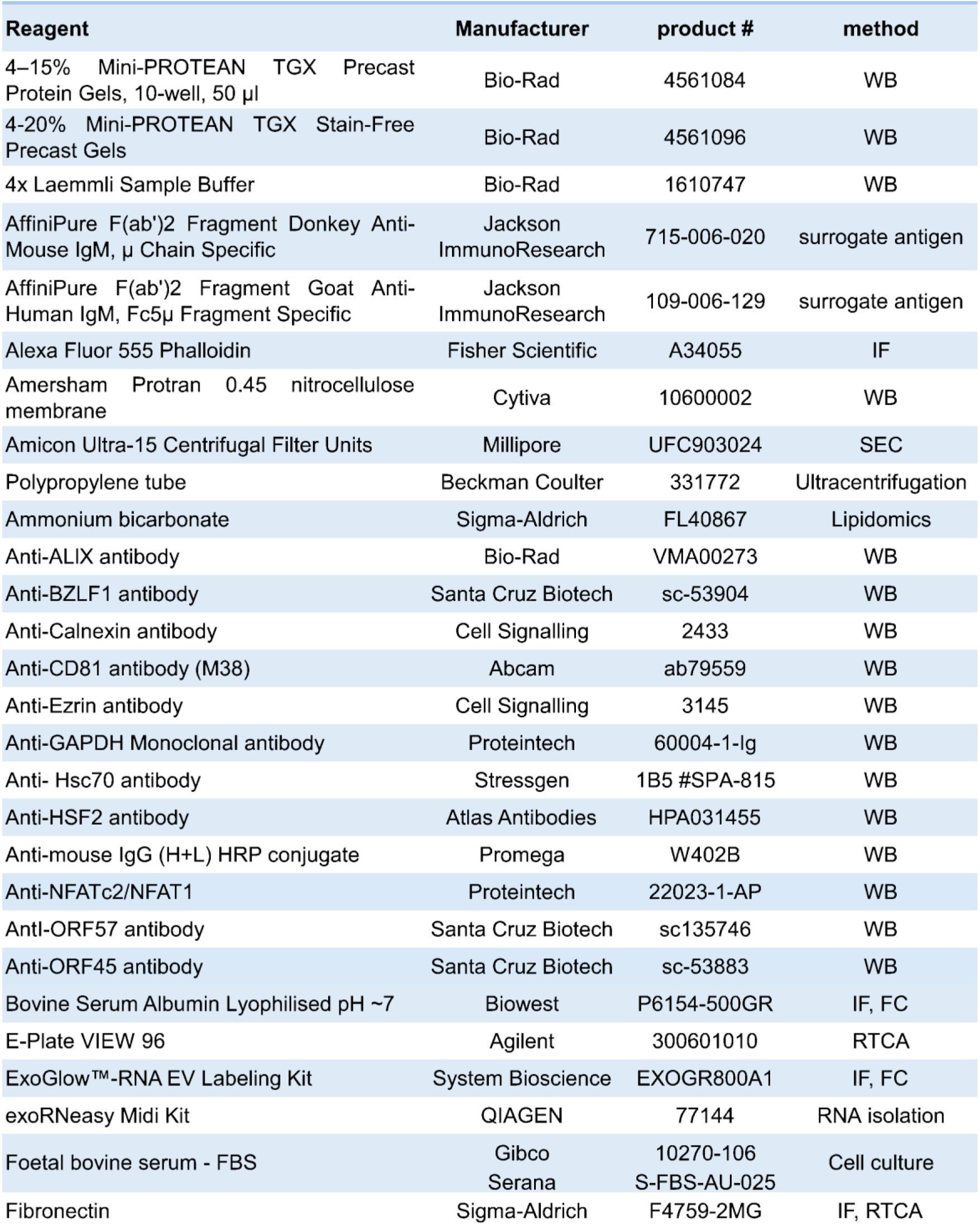
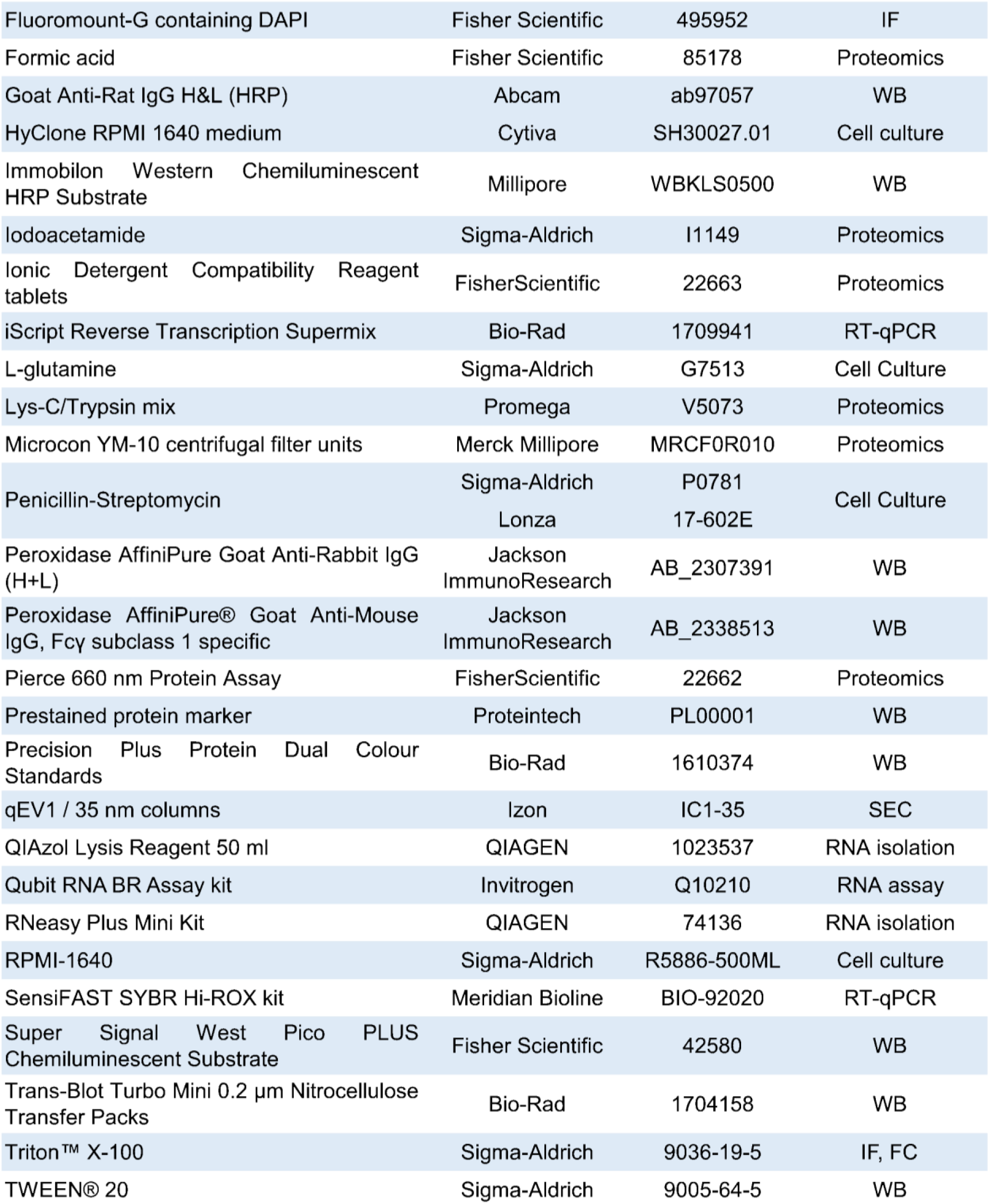
Reagents used in this study with source and application.

## Data Availability

The mass spectrometry raw data have been deposited to the ProteomeXchange Consortium via the PRIDE ^70^ partner repository with the dataset identifier PXD074089. Raw and processed RNA-seq data have been submitted to NCBI GEO database and will be publicly available upon completion of the GEO curation process (https://www.ncbi.nlm.nih.gov/geo/).

## Author Contributions

AM, SH, PKM and SG designed and conceptualized the research; AM, HD, DC, SH, SV, MM, KC, KM performed research, analyzed data, and prepared figures; PKM, SG, MJ provided resources and obtained funding; and AM, SG and PKM wrote the paper with input and review from all authors.

## Supplementary data

**Supplementary Figure S1.**
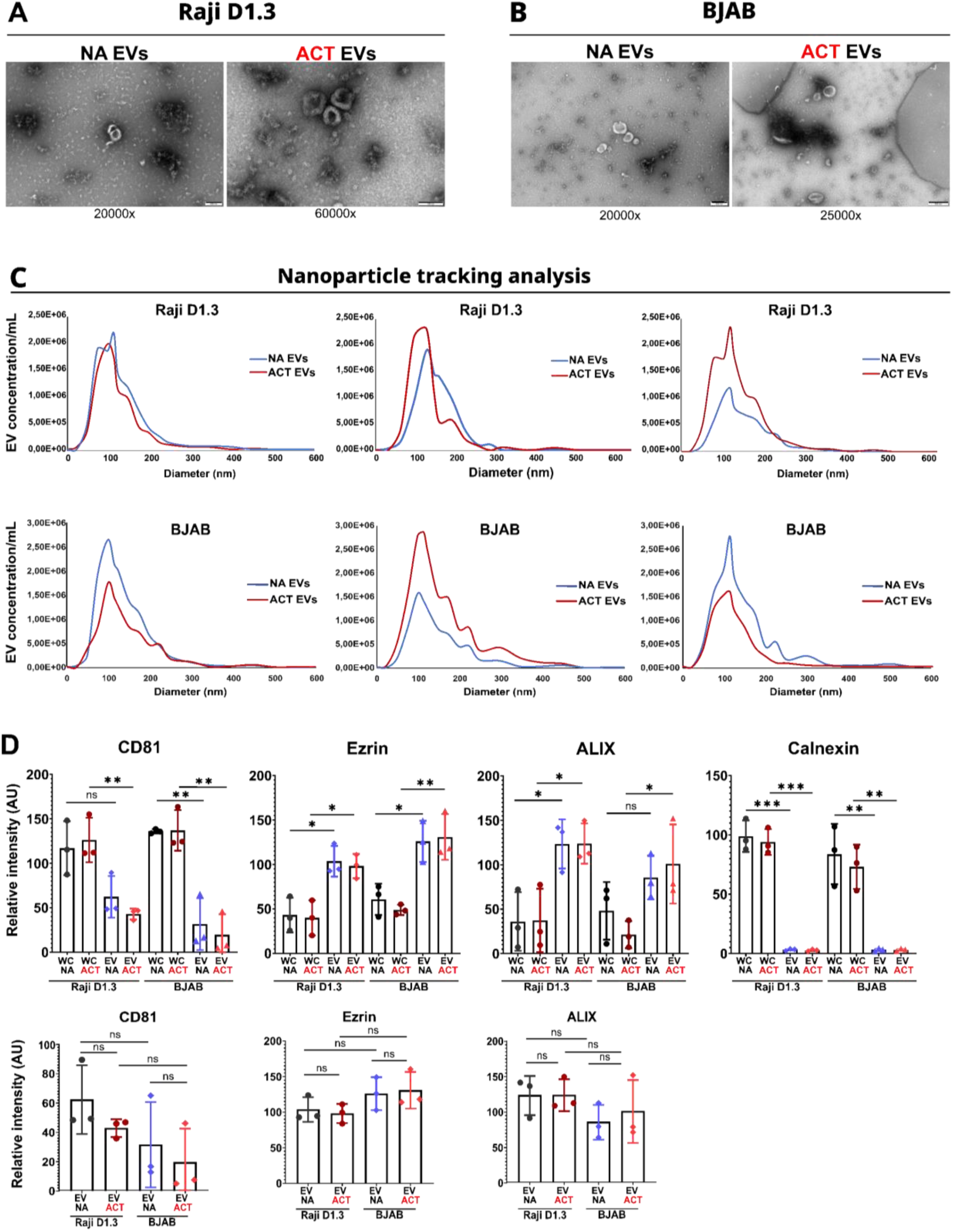
EV characterization. **(A-B)** Raw TEM images from Fig 1.B, zoomed out with magnification. Scale bar 200 nm in all images except for in Raji D1.3 ACT EVs 100 nm. **(C)** Nanoparticle tracking analysis (NTA) plots from three independent experiments, showing EV size distribution profiles of NA and ACT samples from Raji D1.3 and BJAB cell lines. **(D)** Quantification of Western blot analysis for EV and WCL markers from 30 min activated (ACT) and non-activated (NA) Raji D1.3 and BJAB cells (related to Fig. 1). Protein band intensities were measured using ImageJ (FIJI) and normalized to the loading control (GAPDH). Data is presented as mean ± SEM from three independent experiments (N = 3). Statistical significance was calculated using a two-tailed unpaired Student’s t-test in GraphPad. *p < 0.05, **p < 0.01, ***p < 0.001.

**Supplementary Figure S2.**
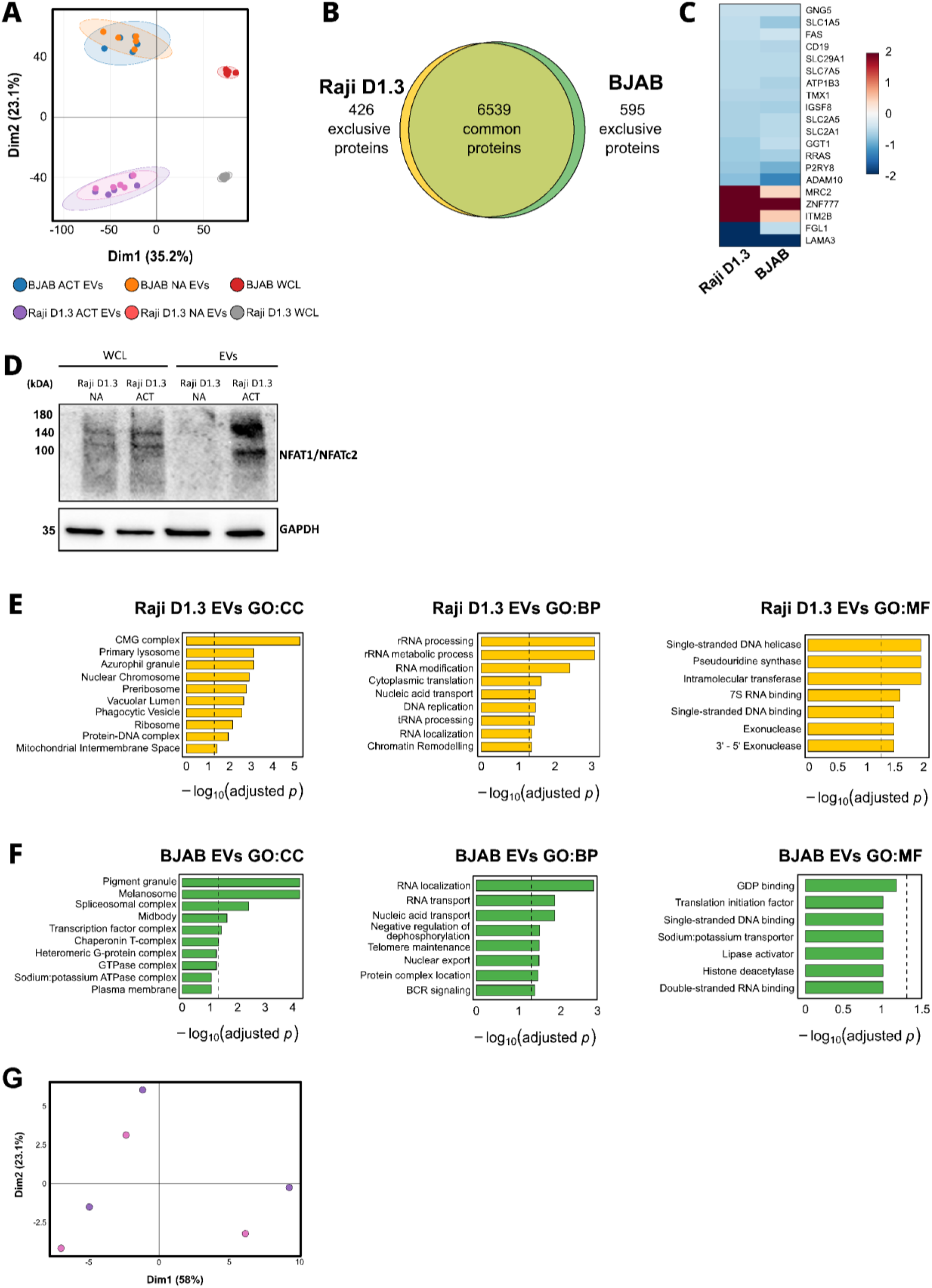
Proteomic and transcriptomic analyses of B-EVs. **(A)** Principal Component Analysis (PCA) of isolated EVs from Act or NA conditions versus WCL from BJAB and Raji D1.3 cell lines. Points represent individual samples (n=5/condition) with 95% confidence ellipses. **(B)** Venn diagram with of proteins identified in BJAB (green) and Raji D1.3 (yellow) EVs showing how many are overlapping and exclusive to each cell line. **(C)** Heatmap of proteins upregulated in Act or NA EVs in both BJAB and Raji D1.3 cell lines (−0.4 < log₂FC > 0.4). Analysis includes differentially expressed. **(D)** Western blot analysis of Raji D1.3 and their corresponding EV samples confirms that NFAT1/NFATc2 protein is enriched only in ACT EV samples, as shown also in the proteomic analysis. **(E, F)** and exclusive proteins; exclusive proteins were assigned an artificial fold change of 2 for visualization. Gene Ontology (GO) enrichment analysis of EV-associated proteins from Raji D1.3 (E) and BJAB (F) cells under ACT or NA conditions. Bar plots show significantly enriched Cellular Component (CC), Biological Process (BP), and Molecular Function (MF) terms (BH-corrected). Bars represent −log₁₀(adjusted p-values); dashed line indicates significance threshold (adjusted p-value < 0.05). (E-F). **(G)** PCA of isolated EVs from Act or NA conditions from Raji D1.3 cell line, transcriptomics analysis. Points represent individual samples (n=3/condition). ACT EVs, purple; NA EVs, pink.

**Supplementary Figure S3.**
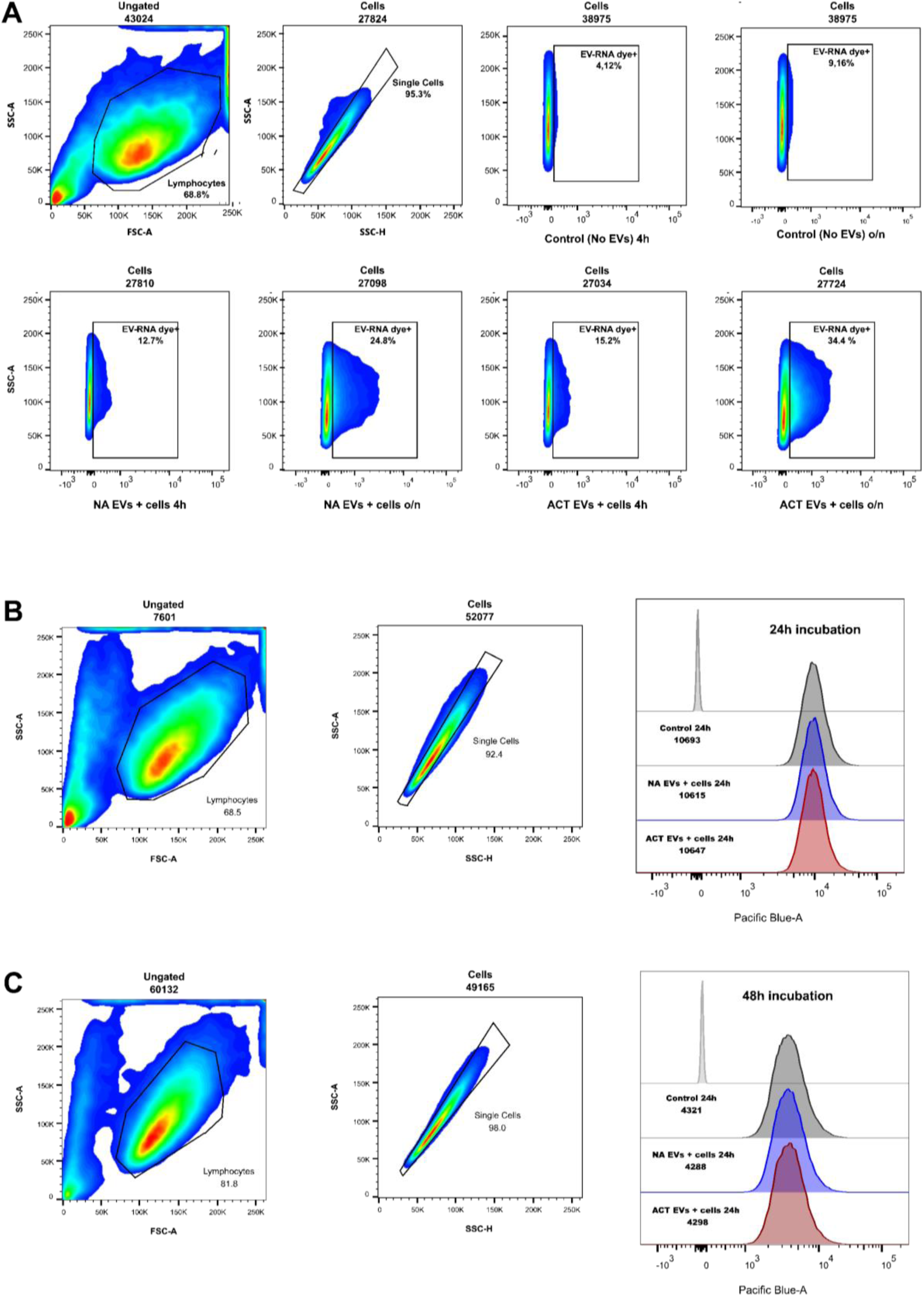
EVs uptake into recipient cells. **(A)** Representative flow cytometry plots from one of four independent experiments (n = 4) showing the percentage of EV-RNA–positive recipient cells and corresponding mean fluorescence intensity after incubation. **(B, C)** Flow cytometry analysis showing no differences in recipient cell proliferation between conditions after 24 or 48 h of incubation (n = 1).

**Supplementary Figure S4.**
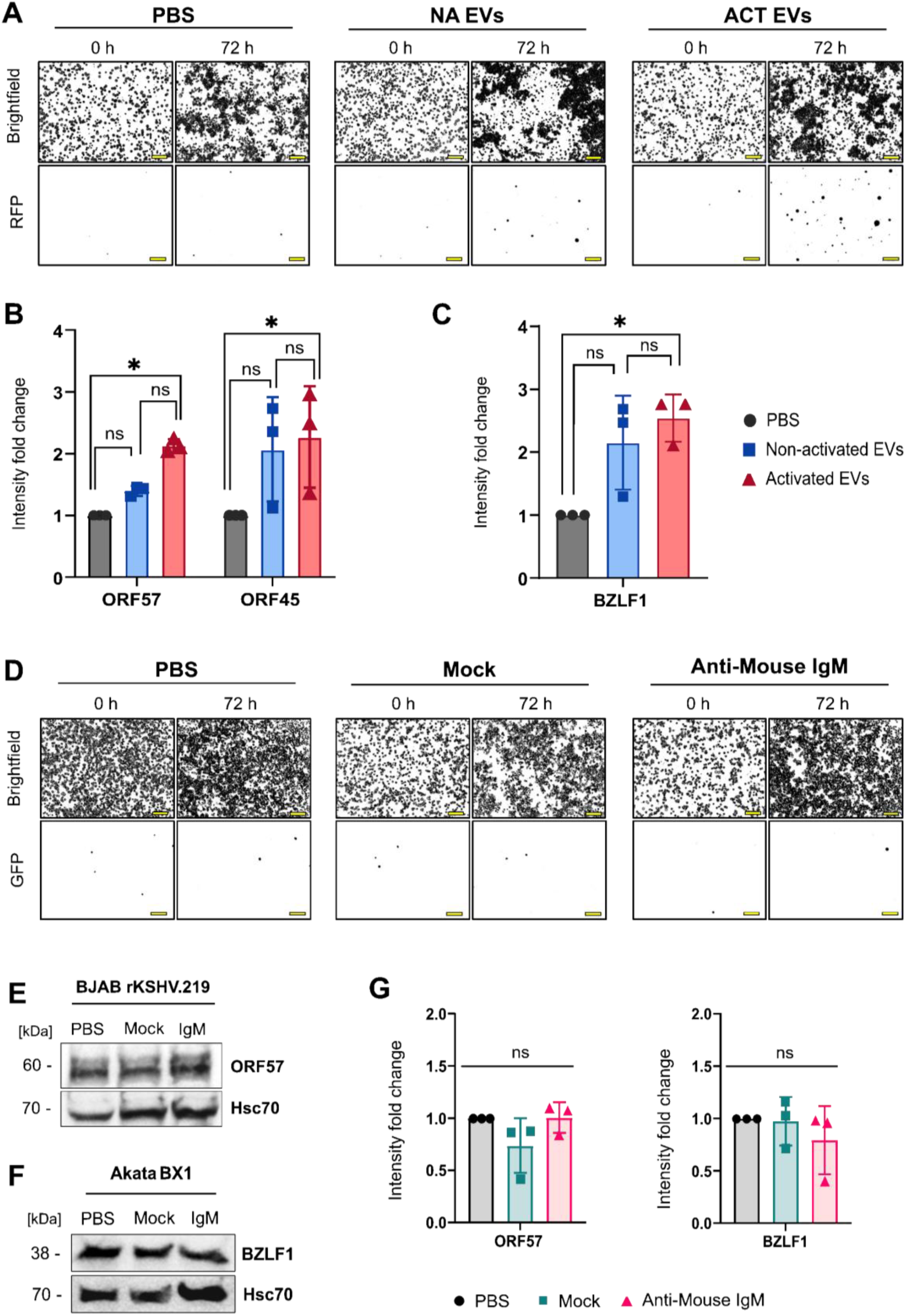
Activated B cell–derived EVs induce EBV and KSHV lytic gene expression. **(A-C)** BJAB rKSHV.219 cells were treated for 72 h with 10 μg of B-EVs isolated from non-activated (NA) or activated (ACT) B cells. **(A)** Representative brightfield (top) and RFP fluorescence (bottom) binary masks at 0 h and 72 h (N = 4). Scale bar, 100 μm. Masks generated in FIJI. **(B)** Quantification of ORF45 and ORF57 immunoblots (Fig. 6), normalized to loading controls (mean ± SEM, N = 3). Two-way ANOVA with Tukey’s multiple-comparisons test. **(C)** Quantification of BZLF1 immunoblots (Fig. 6), normalized to loading controls (mean ± SEM, N = 3). One-way ANOVA. D-F) BJAB rKSHV.219 and EBV-positive Akata BX1 cells were treated for 72 h with surrogate antigen (anti-mouse IgM) alone or supplemented with cell-free medium processed for EV isolation (“Mock”). **(D)** Representative brightfield and RFP binary masks of BJAB rKSHV.219 cells at 0 h and 72 h (N = 4). Scale bar, 100 μm. **(E)** Representative ORF57 immunoblot in BJAB rKSHV.219 cells. **(F)** Representative BZLF1 immunoblot in Akata BX1 cells. Hsc70 served as loading control (N = 3). **(G**) Quantification of ORF57 and BZLF1 immunoblots (panels E and F), normalized to loading controls (mean ± SEM, N = 3). Two-way ANOVA with Tukey’s multiple-comparisons test. *p < 0.05, **p < 0.01, ***p < 0.001, n.s.: non-significant.

**Supplementary Table SI:** Lipidomic analysis, full dataset. Mol% values of lipid species identified in EVs and whole cell (WC) lysates used in comparative analyses.

**Supplementary Table SII:** Proteomic analysis, full dataset. **(A)** List of identified EV proteins in Raji D1.3 parental cell line with > 1 unique peptides (3 > valid values in both resting and activated Evs). **(B)** List of identified EV proteins in BJAB parental cell line with > 1 unique peptides (3 > valid values in both resting and activated Evs). **(C)** List of exclusively identified proteins in either Raji D1.3 parental cell line or EVs with > 1 unique peptides (< 3 valid values). **(D)** List of exclusively identified proteins in either BJAB parental cell line or EVs with > 1 unique peptides (< 3 valid values). **(E)** Output from Spectronaut when using Donkey Proteome as Reference.

## References

1. Busatto, S. et al. Tangential Flow Filtration for Highly Efficient Concentration of Extracellular Vesicles from Large Volumes of Fluid. Cells 7, 273 (2018).

2. Bonsergent, E. et al. Quantitative characterization of extracellular vesicle uptake and content delivery within mammalian cells. Nat. Commun. 12, 1864 (2021).

3. Bonner, S. E. & Willms, E. Intercellular communication through extracellular vesicles in cancer and evolutionary biology. Prog. Biophys. Mol. Biol. 165, 80–87 (2021).

4. Kumar, M. A. et al. Extracellular vesicles as tools and targets in therapy for diseases. Signal Transduct. Target. Ther. 9, 1–41 (2024).

5. Hoshino, A. et al. Extracellular Vesicle and Particle Biomarkers Define Multiple Human Cancers. Cell 182, 1044–1061.e18 (2020).

6. Fanny A. Pelissier Vatter et al. Extracellular vesicle- and particle-mediated communication shapes innate and adaptive immune responses. J. Exp. Med. 218, (2021).

7. Lu, M. et al. The Role of Extracellular Vesicles in the Pathogenesis and Treatment of Autoimmune Disorders. Front. Immunol. 12, (2021).

8. Fariba Mahmoudi & Parichehr Hanachi. Extracellular vesicles of immune cells; immunomodulatory impacts and therapeutic potentials. 248, 109237–109237 (2023).

9. Yang, P. et al. Immune Cell-Derived Extracellular Vesicles – New Strategies in Cancer Immunotherapy. Front. Immunol. 12, 771551 (2021).

10. Hansen, A. S., et al. T-cell derived extracellular vesicles prime macrophages for improved STING based cancer immunotherapy. J. Extracell. Vesicles 12, e12350 (2023).

11. Torralba, D. et al. Priming of dendritic cells by DNA-containing extracellular vesicles from activated T cells through antigen-driven contacts. Nat. Commun. 9, 2658 (2018).

12. Mittelbrunn, M. et al. Unidirectional transfer of microRNA-loaded exosomes from T cells to antigen-presenting cells. Nat. Commun. 2, 282 (2011).

13. Choudhuri, K. et al. Polarized release of T-cell-receptor-enriched microvesicles at the immunological synapse. Nature 507, 118–123 (2014).

14. Raposo, G. et al. B lymphocytes secrete antigen-presenting vesicles. J. Exp. Med. 183, 1161–1172 (1996).

15. Rival, C. et al. B cells secrete functional antigen-specific IgG antibodies on extracellular vesicles. Sci. Rep. 14, 16970 (2024).

16. Buzas, E. I. The roles of extracellular vesicles in the immune system. Nat. Rev. Immunol. 23, 236–250 (2023).

17. Gutknecht, M. F., Holodick, N. E. & Rothstein, T. L. B cell extracellular vesicles contain monomeric IgM that binds antigen and enters target cells. iScience 26, 107526 (2023).

18. Muntasell, A., Berger, A. C. & Roche, P. A. T cell-induced secretion of MHC class II–peptide complexes on B cell exosomes. EMBO J. 26, 4263–4272 (2007).

19. Qazi, K. R., Gehrmann, U., Domange Jordö, E., Karlsson, M. C. I. & Gabrielsson, S. Antigen-loaded exosomes alone induce Th1-type memory through a B cell–dependent mechanism. Blood 113, 2673–2683 (2009).

20. Phan, H.-D. et al. CD24 and IgM Stimulation of B Cells Triggers Transfer of Functional B Cell Receptor to B Cell Recipients Via Extracellular Vesicles. J. Immunol. 207, 3004–3015 (2021).

21. Saunderson, S. C. et al. Induction of Exosome Release in Primary B Cells Stimulated via CD40 and the IL-4 Receptor1. J. Immunol. 180, 8146–8152 (2008).

22. Kuokkanen, E., Šuštar, V. & Mattila, P. K. Molecular control of B cell activation and immunological synapse formation. Traffic Cph. Den. 16, 311–326 (2015).

23. Cyster, J. G. & Allen, C. D. C. B Cell Responses: Cell Interaction Dynamics and Decisions. Cell 177, 524–540 (2019).

24. Batista, F. D. & Harwood, N. E. The who, how and where of antigen presentation to B cells. Nat. Rev. Immunol. 9, 15–27 (2009).

25. Münz, C. Modulation of Epstein-Barr-Virus (EBV)-Associated Cancers by Co-Infections. Cancers 15, 5739 (2023).

26. Li, S., Bai, L., Dong, J., Sun, R. & Lan, K. Kaposi’s Sarcoma-Associated Herpesvirus: Epidemiology and Molecular Biology. in Infectious Agents Associated Cancers: Epidemiology and Molecular Biology (eds Cai, Q., Yuan, Z. & Lan, K.) 91–127 (Springer, Singapore, 2017). doi:10.1007/978-981-10-5765-6_7.

27. Lieberman, P. M. Epigenetics and Genetics of Viral Latency. Cell Host Microbe 19, 619–628 (2016).

28. Gramolelli, S. & Schulz, T. F. The role of Kaposi sarcoma-associated herpesvirus in the pathogenesis of Kaposi sarcoma. J. Pathol. 235, 368–380 (2015).

29. Gramolelli, S. & Ojala, P. M. Kaposi’s sarcoma herpesvirus-induced endothelial cell reprogramming supports viral persistence and contributes to Kaposi’s sarcoma tumorigenesis. Curr. Opin. Virol. 26, 156–162 (2017).

30. Dotto-Maurel, A., Arzul, I., Morga, B. & Chevignon, G. Herpesviruses: overview of systematics, genomic complexity and life cycle. Virol. J. 22, 155 (2025).

31. Kati, S. et al. Activation of the B Cell Antigen Receptor Triggers Reactivation of Latent Kaposi’s Sarcoma-Associated Herpesvirus in B Cells. J. Virol. 87, 8004–8016 (2013).

32. Chen, W., Xie, Y., Li, F., Wen, P. & Wang, L. EBV + B cell-derived exosomes promote EBV-associated T/NK-cell lymphoproliferative disease immune evasion by STAT3/IL-10/PD-L1 pathway. Immunol. Res. 72, 1327–1336 (2024).

33. Takada, K. Cross-linking of cell surface immunoglobulins induces epstein-barr virus in burkitt lymphoma lines. Int. J. Cancer 33, 27–32 (1984).

34. Zhang, K., Lv, D.-W. & Li, R. B Cell Receptor Activation and Chemical Induction Trigger Caspase-Mediated Cleavage of PIAS1 to Facilitate Epstein-Barr Virus Reactivation. Cell Rep. 21, 3445–3457 (2017).

35. Minimal information for studies of extracellular vesicles (MISEV2023): From basic to advanced approaches - Welsh - 2024 - Journal of Extracellular Vesicles - Wiley Online Library. https://isevjournals.onlinelibrary.wiley.com/doi/10.1002/jev2.12404.

36. Konoshenko, M. Yu., Lekchnov, E. A., Vlassov, A. V. & Laktionov, P. P. Isolation of Extracellular Vesicles: General Methodologies and Latest Trends. BioMed Res. Int. 2018, 8545347 (2018).

37. Nielsen, I. Ø. et al. Comprehensive Evaluation of a Quantitative Shotgun Lipidomics Platform for Mammalian Sample Analysis on a High-Resolution Mass Spectrometer. J. Am. Soc. Mass Spectrom. 31, 894–907 (2020).

38. van Meer, G., Voelker, D. R. & Feigenson, G. W. Membrane lipids: where they are and how they behave. Nat. Rev. Mol. Cell Biol. 9, 112–124 (2008).

39. Sokoya, T. et al. Pathogenic variants of sphingomyelin synthase SMS2 disrupt lipid landscapes in the secretory pathway. eLife 11, e79278 (2022).

40. Gupta, N. & DeFranco, A. L. Lipid rafts and B cell signaling. Semin. Cell Dev. Biol. 18, 616–626 (2007).

41. Peeters, R. & Jellusova, J. Lipid metabolism in B cell biology. Mol. Oncol. 18, 1795–1813 (2024).

42. Pathan, M. et al. Vesiclepedia 2019: a compendium of RNA, proteins, lipids and metabolites in extracellular vesicles. Nucleic Acids Res. 47, D516–D519 (2019).

43. Mathivanan, S. & Simpson, R. J. ExoCarta: A compendium of exosomal proteins and RNA. PROTEOMICS 9, 4997–5000 (2009).

44. Garcia-Martin, R., Brandao, B. B., Thomou, T., Altindis, E. & Kahn, C. R. Tissue differences in the exosomal/small extracellular vesicle proteome and their potential as indicators of altered tissue metabolism. Cell Rep. 38, 110277 (2022).

45. Vagner, T. et al. Protein Composition Reflects Extracellular Vesicle Heterogeneity. PROTEOMICS 19, 1800167 (2019).

46. Hurwitz, S. N. et al. Proteomic profiling of NCI-60 extracellular vesicles uncovers common protein cargo and cancer type-specific biomarkers. Oncotarget 7, 86999–87015 (2016).

47. Peng, S. L., Gerth, A. J., Ranger, A. M. & Glimcher, L. H. NFATc1 and NFATc2 Together Control Both T and B Cell Activation and Differentiation. Immunity 14, 13–20 (2001).

48. Vercellino, I. & Sazanov, L. A. The assembly, regulation and function of the mitochondrial respiratory chain. Nat. Rev. Mol. Cell Biol. 23, 141–161 (2022).

49. Cohen, J. I. Herpesvirus latency. J. Clin. Invest. 130, 3361–3369 (2020).

50. Laichalk, L. L. & Thorley-Lawson, D. A. Terminal Differentiation into Plasma Cells Initiates the Replicative Cycle of Epstein-Barr Virus In Vivo. J. Virol. 79, 1296–1307 (2005).

51. Myoung, J. & Ganem, D. Generation of a doxycycline-inducible KSHV producer cell line of endothelial origin: maintenance of tight latency with efficient reactivation upon induction. J. Virol. Methods 174, 12–21 (2011).

52. Fields, B. N. Fields Virology. (Wolters Kluwer Health/Lippincott Williams & Wilkins, Philadelphia, 2013).

53. Lin, K.-M., Weng, L.-F., Chen, S.-Y. J., Lin, S.-J. & Tsai, C.-H. Upregulation of IQGAP2 by EBV transactivator Rta and its influence on EBV life cycle. J. Virol. 97, e00540–23 (2023).

54. Cutrone, L. et al. Heat shock factor 2 regulates oncogenic gamma-herpesvirus gene expression by remodeling the chromatin at the ORF50 and BZLF1 promoter. PLOS Pathog. 21, e1013108 (2025).

55. Biswas, A. et al. Inhibition of polo-like kinase 1 (PLK1) facilitates reactivation of gamma-herpesviruses and their elimination. PLOS Pathog. 17, e1009764 (2021).

56. Zhang, H. et al. Mitochondrial protein, TBRG4, modulates KSHV and EBV reactivation from latency. PLOS Pathog. 18, e1010990 (2022).

57. Ayre, D. C. et al. CD24 induces changes to the surface receptors of B cell microvesicles with variable effects on their RNA and protein cargo. Sci. Rep. 7, 8642 (2017).

58. Phan, H.-D., et al. CD24 Regulates the Formation of Ectosomes in B Lymphocytes. J. Extracell. Vesicles 14, e70093 (2025).

59. Ayre, D. C. et al. Dynamic regulation of CD24 expression and release of CD24-containing microvesicles in immature B cells in response to CD24 engagement. Immunology 146, 217–233 (2015).

60. Flemington, E. & Speck, S. H. Autoregulation of Epstein-Barr virus putative lytic switch gene BZLF1. J. Virol. 64, 1227–1232 (1990).

61. Xie, J., Ajibade, A. O., Ye, F., Kuhne, K. & Gao, S.-J. Reactivation of Kaposi’s sarcoma-associated herpesvirus from latency requires MEK/ERK, JNK and p38 multiple mitogen-activated protein kinase pathways. Virology 371, 139–154 (2008).

62. Brambilla, L., Boneschi, V., Zampieri, M., Bruognolo, L. & Fossati, S. Persistently Recurring Mediterranean Kaposi’s Sarcoma on Skin Grafts. Int. J. Dermatol. 35, 362–364 (1996).

63. Niedt, G. W. & Prioleau, P. G. Kaposi’s sarcoma occurring in a dermatome previously involved by herpes zoster. J. Am. Acad. Dermatol. 18, 448–451 (1988).

64. Bringeland, E. A. et al. Epstein–Barr Virus and Clinico-Endoscopic Characteristics of Gastric Remnant Cancers Compared to Proximal Non-Remnant Cancers: A Population-Based Study. Cancers 16, 2000 (2024).

65. Klein, G. et al. Continuous Lymphoid Cell Lines with Characteristics of B Cells (Bone-Marrow-Derived), Lacking the Epstein-Barr Virus Genome and Derived from Three Human Lymphomas. Proc. Natl. Acad. Sci. 71, 3283–3286 (1974).

66. Vieira, J. & O’Hearn, P. M. Use of the red fluorescent protein as a marker of Kaposi’s sarcoma-associated herpesvirus lytic gene expression. Virology 325, 225–240 (2004).

67. Herzog, R. et al. LipidXplorer: A Software for Consensual Cross-Platform Lipidomics. PLOS ONE 7, e29851 (2012).

68. Wiśniewski, J. R., Zougman, A., Nagaraj, N. & Mann, M. Universal sample preparation method for proteome analysis. Nat. Methods 6, 359–362 (2009).

69. Callister, S. J. et al. Normalization Approaches for Removing Systematic Biases Associated with Mass Spectrometry and Label-Free Proteomics. J. Proteome Res. 5, 277–286 (2006).

70. Perez-Riverol, Y. et al. The PRIDE database resources in 2022: a hub for mass spectrometry-based proteomics evidences. Nucleic Acids Res. 50, D543–D552 (2022).

